# Lower hemoglobin levels associate with lower body mass index and healthier metabolic profile

**DOI:** 10.1101/472142

**Authors:** Juha Auvinen, Joona Tapio, Ville Karhunen, Johannes Kettunen, Raisa Serpi, Elitsa Y. Dimova, Pasi Soininen, Tuija Tammelin, Juha Mykkänen, Katri Puukka, Mika Kähönen, Emma Raitoharju, Terho Lehtimäki, Mika Ala-Korpela, Olli T. Raitakari, Sirkka Keinänen-Kiukaanniemi, Marjo-Riitta Järvelin, Peppi Koivunen

## Abstract

Hemoglobin (Hb) is the main carrier of oxygen. In general, high-end Hb levels within the normal range are considered beneficial for health^1^. However, activation of the hypoxia response has been shown to protect mice against metabolic dysfunction^2-4^. We used Hb levels as an indicator for oxygenation status and studied its association with >170 anthropometric and metabolic parameters in two Finnish birth cohorts both in cross-sectional and longitudinal design (max n = 7,175). Here we show a positive linear association between Hb levels and body mass index (BMI). Subjects with the lower Hb levels had better glucose tolerance, lower cholesterol and blood pressure levels, less adverse metabolite profiles and lower inflammatory load. Notably, these associations were not only mediated by the lower BMI, and the effect size of many of them increased with age. Polygenic risk score (PRS) analyses indicated shared genetic determinants between Hb levels and BMI, insulin, triglyceride and HDL cholesterol levels. Mendelian randomization (MR) analyses could not demonstrate causal relationships between Hb and metabolic parameters. However, manipulation of Hb levels by venesection in mice showed evidence for causal associations with body weight and metabolic parameters. Our findings suggest that lower-end normal Hb levels may be favorable for systemic metabolism involving mild chronic activation of the hypoxia response. Therefore modulation of Hb levels could be a novel strategy towards maintenance of metabolic health.

---

Hb, an iron-containing metalloprotein in red blood cells, is the main carrier of oxygen. Its levels directly affect arterial oxygen concentration and thereby tissue oxygenation^5, 6^. Hb levels are regulated genetically and environmentally, and they vary by sex, race, age and altitude^1, 7^. Individual’s Hb levels during adult life are, however, very stable and used *e*.*g*. in the athlete’s biological passport to detect doping^8^. When tissues encounter reduced oxygen levels as a key transcriptional response the hypoxia-inducible factor (HIF) becomes stabilized. It upregulates genes that induce oxygen delivery and reduce its usage, such as those regulating energy metabolism^9^. The stability of HIF is governed by HIF prolyl 4-hydroxylases (HIF-P4Hs), which require oxygen for catalysis and targeting HIF for destruction^9^. Recent studies show that inhibition of HIF-P4Hs, which stabilizes HIF in normoxia, protects mice from obesity, metabolic dysfunction and associated diseases^2-4, 10, 11^. To investigate oxygenation status with regards to metabolic health in human population at a larger scale, we explored the possibility to use the normal variation in Hb levels as a surrogate measure. To test this we first evaluated the association of Hb levels with key metabolic parameters in C57Bl/6 mice. We found positive associations between Hb levels and body weight, less favorable glucose tolerance and homeostatic model assessment-insulin resistance (HOMA-IR) scores, respectively (Figs. 1a-1c), suggesting that Hb levels may influence metabolic health.

**Figure 1.**
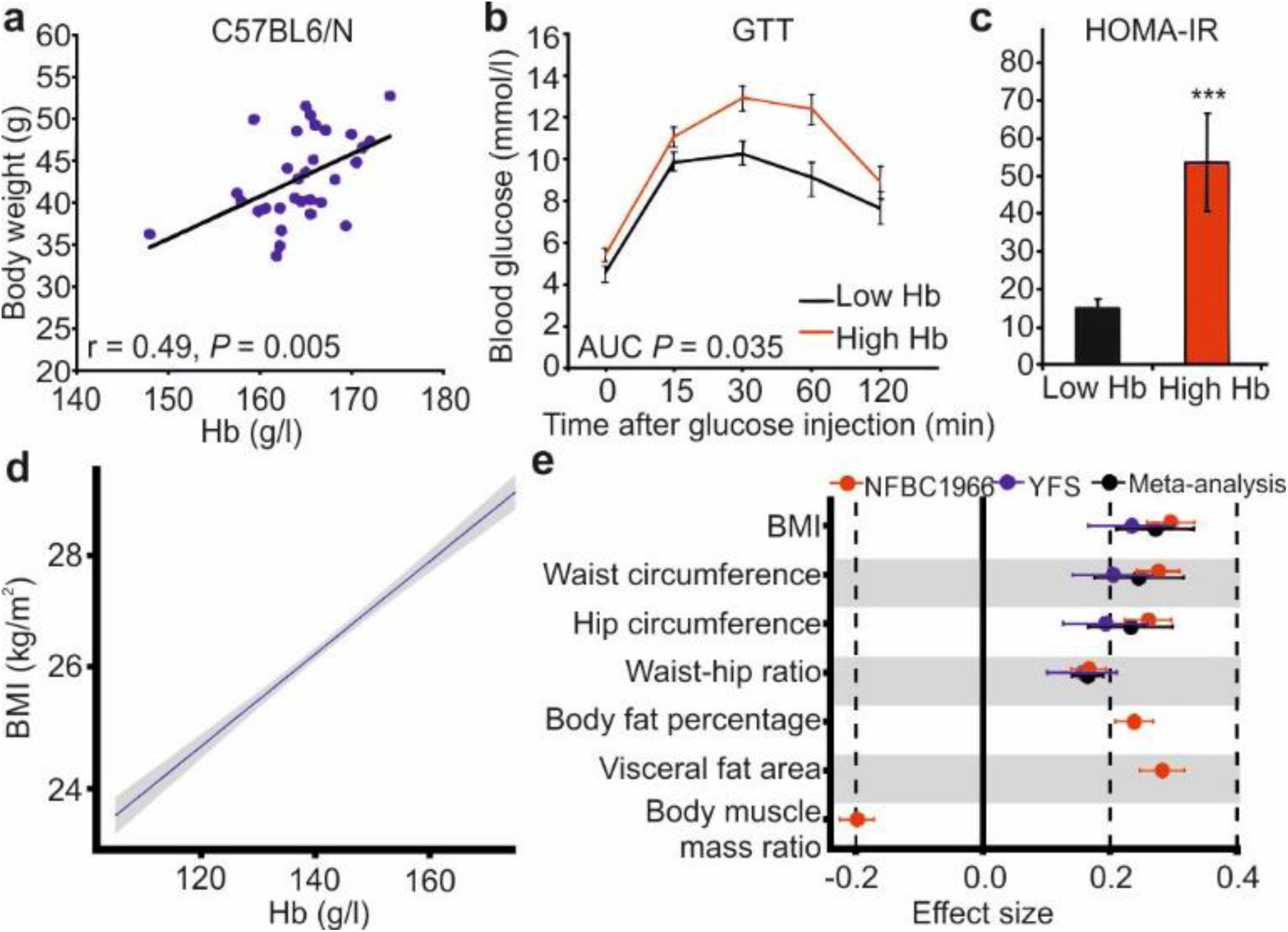
Association of Hb levels with anthropometric and metabolic measures in mice and human. **a**, Association of Hb levels with body weight in 1-year-old C57Bl/6 male mice (n = 31). **b,** Association of the highest quartile (High Hb) and the lowest quartile (Low Hb) of Hb levels with glucose tolerance test (GTT) area under the curve (AUC) (**b**) and HOMA-IR scores, respectively, in the 1-year-old C57Bl/6 male mice (**c**). **d,** Association of Hb levels with BMI in NFBC1966 at age of 46. Unadjusted regression line (blue) with 95% confidence intervals (gray) of blood Hb levels (g/l) with BMI (kg/m^2^). **e,** Forest plot representing of the effect size of association of Hb levels with log(BMI), waist and hip circumference and waist-hip ratio in NFBC1966 (red), YFS (blue) and meta-analysis (black), respectively, and Hb levels with body fat percentage, visceral fat area and body muscle mass ratio in NFBC1966, respectively. Effects sizes are reported in SD units. The data in **a** is average ± SEM. \*\*\**P* <0.001. The data in **b** was adjusted sex, smoking and physical activity.

We then examined association of Hb levels in humans with anthropometric and metabolic parameters in cross-sectional and longitudinal design from 31 to 46 years in the population based, sea-level Northern Finland Birth Cohort 1966 (NFBC1966)^12, 13^ (n = 5,351), and searched for a replication in another sea-level cohort, Cardiovascular Risk in Young Finns Study (YFS)^14^ (n = 1,824) at 42 years. The statistical analyses were adjusted for potential confounding factors, and as appropriate, for BMI (see Methods and Extended data Figure 1 and Table 1). We found positive linear association between Hb levels and BMI in NFBC1966 at age of 46 (Fig. 1d, effect size, log-transformed 0.296, *p* = 4.73 × 10^-58^, Extended data Fig. 2a), and this association replicated in YFS at 42 years (Fig. 1e, Extended data Table 2). The increase of 10 g/l in Hb corresponded to an increase of 3.5% in BMI (Fig. 1b, Extended data Tables 2 and 3). Hb levels were also positively associated with waist circumference, hip circumference, waist-hip ratio, body fat percentage and visceral fat area, and negatively with body muscle percentage (Fig. 1b, Extended data Tables 2 and 3). Thus the subjects with the lower Hb levels appeared less abdominally obese and having more muscle tissue. We found similar associations for hematocrit levels and red blood cell counts with BMI as for Hb (Extended data Fig. 2 and Table 4).

We next showed by associating Hb and fasting glucose and insulin levels, and the derived insulin resistance, sensitivity and beta cell function indices (HOMA-IR, HOMA-B, Matsuda^15^) from an oral glucose tolerance test that the subjects with lower Hb levels had better glucose tolerance and insulin sensitivity than those with higher Hb levels (Fig. 2a, Extended data Table 5). There was also evidence for positive association between Hb and levels of systolic and diastolic blood pressure, serum total cholesterol, LDL cholesterol, triglycerides and serum high sensitive C-reactive protein (CRP), respectively, and negative association between Hb and HDL cholesterol levels (Fig. 2a, Extended data Table 5). The effect sizes of these associations weakened when additionally adjusted for BMI, but all except that for CRP remained statistically significant (P < 0.02, Fig. 2a, Extended data Table 5).

**Figure 2.**
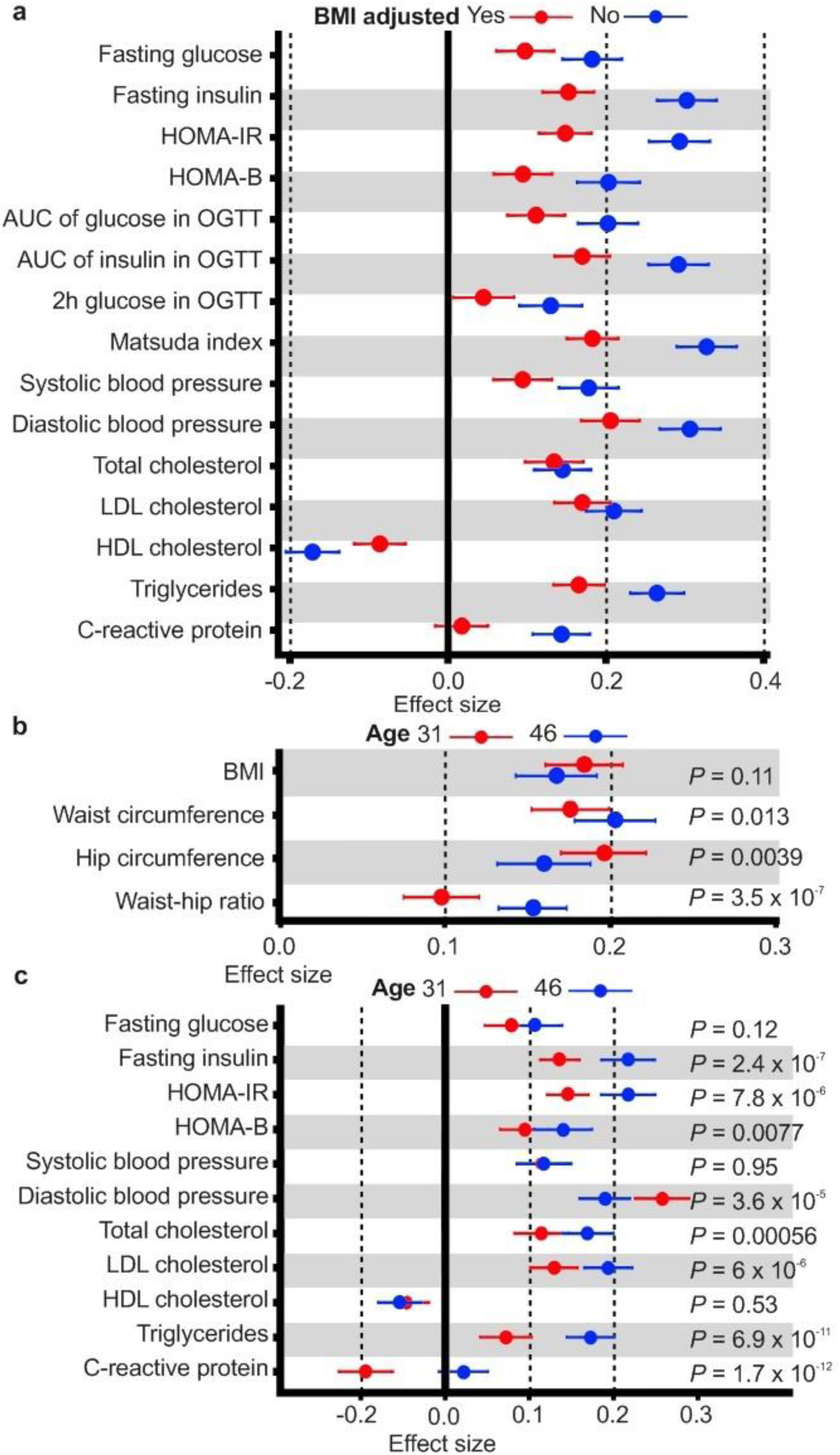
Hb levels associate with key metabolic parameters. Forest plots representing (**a**) the effect sizes of the association in SD units of Hb levels with fasting glucose and log(fasting insulin) levels, log(HOMA-IR) and log(HOMA-B) indexes, log(area under the curve (AUC) of glucose in oral glucose tolerance test (OGTT))*, log(AUC of insulin in OGTT)* and 2 h glucose in levels in a 2 h OGTT*, log(Matsuda index)*, systolic and diastolic blood pressure, fasting serum cholesterol, LDL cholesterol, HDL cholesterol and log(triglyceride) and log(high-sensitivity C-reactive protein (CRP)) levels in meta-analysis of NFBC1966 at age of 46 and YFS at mean age of 42 years, respectively. Asterisk indicates the associations only analyzed in NFBC1966. The data were adjusted sex, smoking and physical activity (blue), and additionally for BMI (red). **b,** Effect sizes of association in SD units of Hb levels with log(BMI), waist and hip circumference and waist-hip ratio in NFBC1966 at age of 31 (red) and 46 (blue) years, respectively. The *P* values represent the difference in effect size between 31 and 46 years. **c**, Effect sizes of the association in SD units of Hb levels with fasting glucose and log(fasting insulin) levels, HOMA-IR and HOMA-B indexes, systolic and diastolic blood pressure, fasting serum cholesterol, LDL cholesterol, HDL cholesterol and triglyceride levels and log(high-sensitivity CRP) in NFBC1966 at age of 31 (red) and 46 (blue) years, respectively. The *P* values for the difference in effect size between age 31 and 46 years are indicated. The data in **b** and **c** were adjusted for sex, smoking and physical activity, and in **c** for BMI.

To evaluate in a longitudinal set up the association of Hb levels with anthropometric and metabolic measures, we compared the effect sizes of these associations in subjects of NFBC1966 that had been examined both at the age of 31 and 46 years (n = 3,624). As expected, the mean Hb levels had stayed unchanged from 31 to 46 years (Extended data Figure 3). In general, the effect sizes of the associations increased with age or stayed the same (Fig. 2b), the most pronounced increases being found for the association of Hb levels with levels of triglycerides and insulin, HOMA-IR and waist-hip-ratio, respectively (Fig. 2c, Extended data Tables 6 and 7).

We then evaluated the association between Hb levels and systemic metabolite profiles of 150 variables analyzed by nuclear magnetic resonance (NMR) metabolomics. Evidence for associations of Hb levels was found with lipoprotein subclass particle concentrations and particle sizes, and levels of ApoB, triglycerides and cholesterol in lipoproteins, fatty acids (FAs), lactate, glycerol, branched-chain amino acids (isoleucine, leucine, valine), aromatic amino acids (phenylalanine, tyrosine), the inflammatory glycoprotein acetyls, creatinine, albumin and ketone bodies, respectively, these associations being mostly positive (Fig. 3, Extended data Table 8). Higher levels of branched-chain and aromatic amino acids, glycoprotein acetyls and lactate, associated with the higher Hb levels here (Fig. 3, Extended data Table 8), have previously been associated with insulin resistance^16^, and the first two with adiposity^17^. Moreover, higher phenylalanine levels and higher ratio of monounsaturated FAs to all FAs, associated with higher Hb levels here (Fig. 3, Extended data Table 8), have earlier been associated with increased risk for cardiovascular events^18^. The magnitudes of cross-sectional and longitudinal effect estimates of the Hb level difference from 31 to 46 years on the NMR metabolite levels in NFBC1966 were consistent (Extended data Figure 4). Furthermore, the estimated slope for regression of longitudinal effects on cross-sectional effects (0.97, 95% CI 0.89 to 1.05) indicated similar effects sizes for both the between-subject and the within-subject effects (Extended data Figure 4). Altogether, lower Hb levels associated with less adverse metabolite profiles.

**Figure 3.**
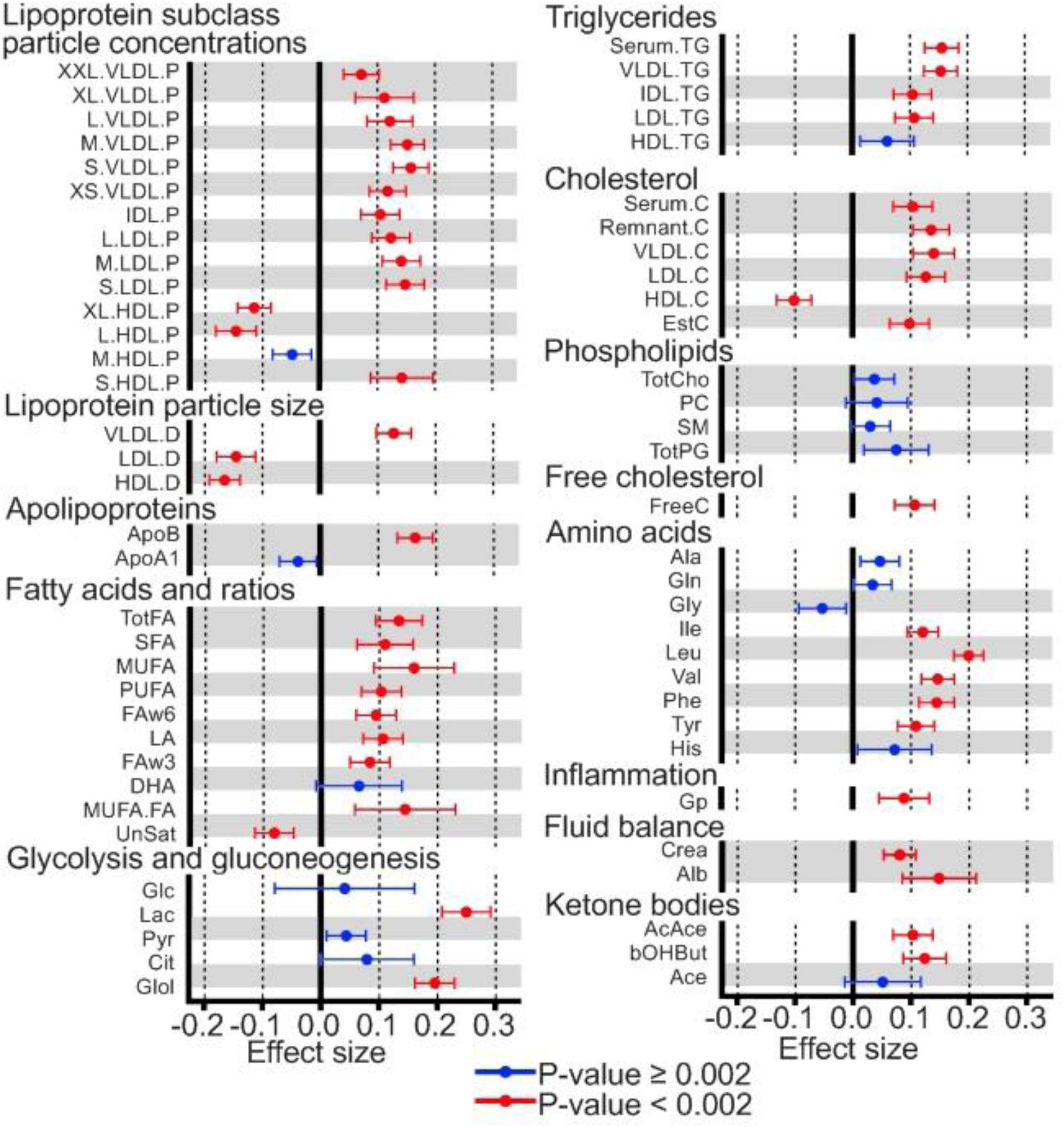
Higher Hb levels associate with metabolic signatures of adiposity, insulin resistance and cardiovascular risk. Effect sizes in SD units of association of Hb levels with systemic metabolite levels in random effects meta-analysis of NFBC1966 at age of 46 and YFS at mean age of 42 years. The data were adjusted for BMI, sex, smoking and physical activity and corrected for multiple testing.

We found evidence for a positive association of Hb levels and oxygen consumption at rest and oxygen consumption and body fat percentage in a subpopulation of NFBC1966^19^ (Fig. 4a, Extended data Fig. 1 and Table 9). Thus the individuals with lower Hb levels, and less fat, consumed less oxygen.

**Figure 4.**
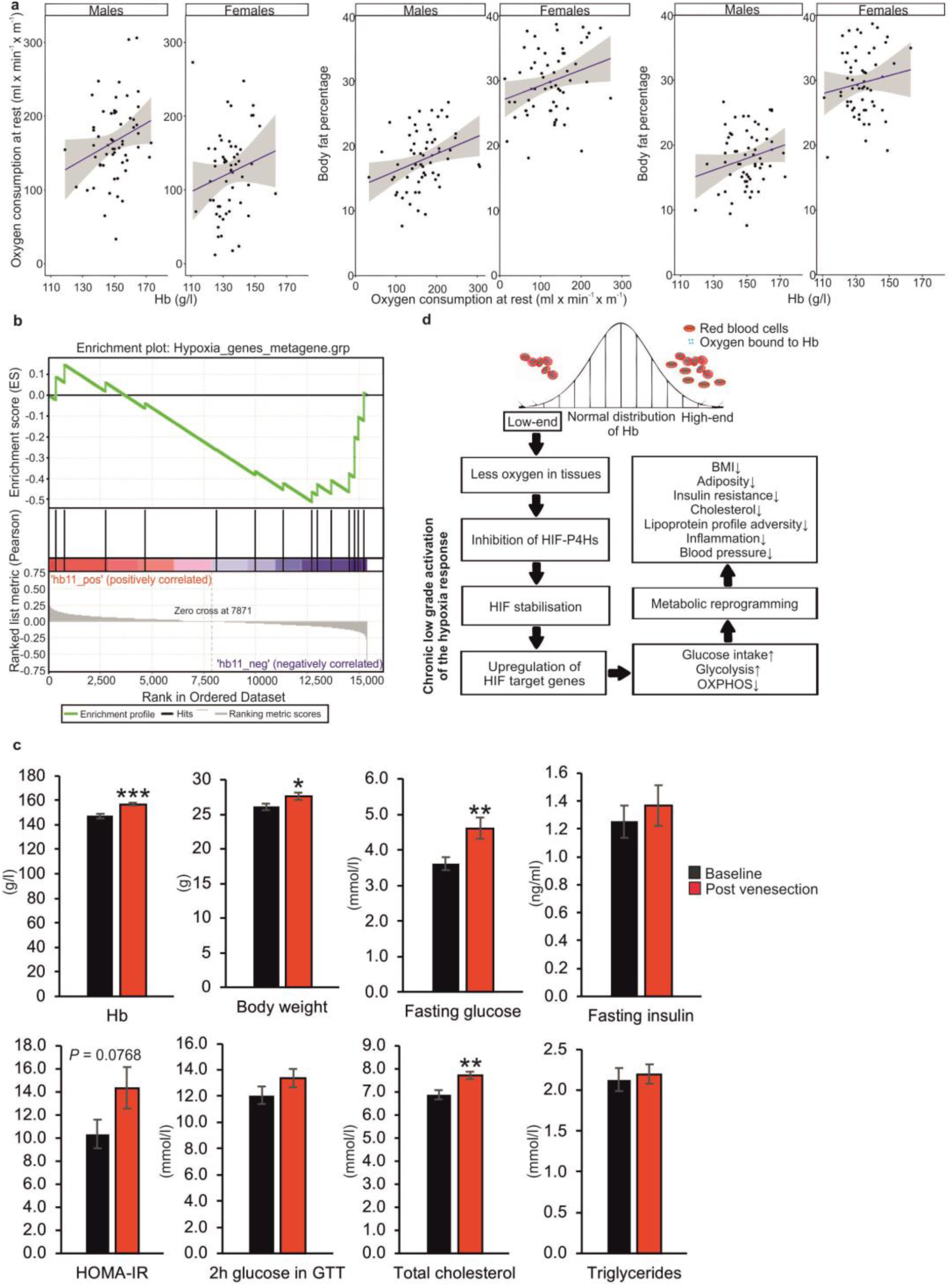
Lower Hb levels associate with decreased oxygen consumption at rest and upregulation of hypoxia-induced genes. **a,** Associations of Hb levels with oxygen consumption at rest (Effect size (Beta) = 0.26, *P* = 0.02 for meta-analysis of both genders), oxygen consumption at rest with body fat percentage (Effect size (Beta) = 0.19, *P* = 0.001 for meta-analysis of both genders) and Hb levels with body fat percentage (Effect size (Beta) = 0.14, *P* = 0.057 for meta-analysis of both genders) in a subpopulation (n = 123) of NFBC1966 at age of 31. **b**, Gene-set enrichment analysis of the YFS lowest Hb quartile (Hb < 132 g/l, n = 392) vs. YFS highest Hb quartile (Hb > 152 g/l, n = 371) and hypoxia-induced genes. Cohort n = 1,636. The analysis was adjusted for sex, age, numbers of thrombocytes and leukocytes, the first five principal components of the transcriptomics data, BMI, research center and three technical microchip variables. **c,** Hb levels, body weight, fasting blood glucose, fasting serum insulin, HOMA-IR, 2h blood glucose levels in GTT, total cholesterol and triglyceride levels of 3-month-old C57Bl/6 male mice (n = 21) before (baseline) and 2 weeks after venesection (post venesection). **d**, Schematic summary of the discovered associations. Hypothesized pathways are depicted by dashed lines. Data in **c** is average ± SEM. \**P* <0.05, \*\**P* <0.01, \*\*\**P* <0.001.

HIF-P4Hs act as cellular oxygen sensors^20^. Even slight reductions in tissue oxygenation, such as those stemming from the normal variation in Hb levels, may result via HIF-P4H inhibition to upregulation of hypoxia-induced genes. To test this hypothesis we analyzed in a subpopulation of YFS^14^ (n=1,636) association of Hb levels with the expression of a set of hypoxia-induced genes (Extended data Table 10) in whole-blood genome wide-expression data. After adjustment for confounding factors, we found evidence for geneset level transcriptional activation of hypoxia-induced genes in individuals in the lowest Hb quartile (Hb < 132 g/l) compared to the highest quartile (Hb > 152 g/l) (Figure 4b).

We examined also the potential causality of Hb levels on BMI and key metabolic measures (and *vice versa*) using bidirectional MR analyses as well as explored shared genetic variation between them using PRS analyses. MR analyses were carried out using the European ancestry summary statistics from genome-wide meta-analysis studies^7, 21^ for these variables. Neither inverse variance weighting (IVW) nor MR-Egger regression method gave clear evidence for causality on any parameter to either direction. PRS analyses gave evidence for shared genetic determinants between Hb and BMI and levels of insulin, triglycerides and HDL cholesterol (Extended data Table 11 and Figures 5 and 6). Thus with the currently available data sets we could not evidence causality for either direction. Its untangling would certainly require a larger Hb genome-wide association material to overcome the weakness of genetic instrument to show potential causality in human data (Extended data Table 12). We therefore went further with the animal experiments and manipulated Hb levels in C57Bl/6 mice by venesection, and showed that in two weeks the induced erythropoiesis increased their Hb levels which associated with enlarged body weight, insulin resistance and cholesterol levels (Fig. 4c, Extended data Figure 7).

While no association of Hb levels with overall metabolic health has been reported before, high Hb or hematocrit levels have earlier been associated in individual epidemiological studies with increased insulin resistance, hypertension, hypercholesterolemia or hypertriglyceridemia^22-25^. Although the underlying mechanisms have been poorly understood hyperviscosity or changes in plasma volume have been suggested as mediators of these associations. The HIF pathway was only discovered after these early analyses. One of the key adjustments in the hypoxia response is transcriptional reprogramming of energy metabolism to less oxygen consuming while less ATP producing, more inefficient mode^9, 26^ (Fig. 4d). The data presented here suggest that the lower Hb levels activated the hypoxia response which could mediate the beneficial effects on metabolic health. Individuals being environmentally exposed to hypoxia by living at high altitude have been reported to have a lower fasting glycemia and better glucose tolerance compared to those at near sea level, and demographic studies associate living at high altitude with protection against obesity and diabetes^27, 28^. Further, clinical trials with HIF-P4H inhibitors for treatment of anemia and ischemia reported ≥20% reduced cholesterol levels in patients receiving these therapeutics^29, 30^. While more mechanistic studies are needed, our human and experimental data show overall that Hb levels seem to contribute to metabolic health and lower Hb levels indicate lower cardiometabolic risk.

## Online Content

Methods, along with any additional Extended Data display items are available in the online version of the paper. References unique to these sections appear only in the online paper.

## Acknowledgements

This study was supported by the Academy of Finland Grants 266719 and 308009, the S. Jusélius Foundation, the Emil Aaltonen Foundation and the Jane and Aatos Erkko Foundation to P.K. NFBC1966 received financial support from University of Oulu Grant no. 65354 and 24000692, Oulu University Hospital Grant no. 2/97, 8/97 and 24301140, Ministry of Health and Social Affairs Grant no. 23/251/97, 160/97, 190/97, National Institute for Health and Welfare, Helsinki Grant no. 54121, Regional Institute of Occupational Health, Oulu, Finland Grant no. 50621, 54231, ERDF European Regional Development Fund Grant no. 539/2010 A31592, the European Union’s Horizon 2020 research and innovation programme grant agreement no. 633595 (DynaHEALTH), no. 733206 (LifeCycle) and no. 643774 (iHEALTH-T2D), Academy of Finland, University Hospital Oulu and NHLBI grant 5R01HL087679-02 through the STAMPEED program. We thank the late professor Paula Rantakallio (launch of NFBC1966), the participants in the 46-year study and the NFBC project center. The Young Finns Study has been financially supported by the Academy of Finland: grants 286284 (T.L.), 134309 (Eye), 126925, 121584, 124282, 129378 (Salve), 117787 (Gendi), and 41071 (Skidi); the Social Insurance Institution of Finland, Competitive State Research Financing of the Expert Responsibility area of Kuopio, Tampere and Turku University Hospitals (grant X51001), Juho Vainio Foundation, Paavo Nurmi Foundation, Finnish Foundation for Cardiovascular Research, Finnish Cultural Foundation, Tampere Tuberculosis Foundation, Emil Aaltonen Foundation, Yrjö Jahnsson Foundation, Signe and Ane Gyllenberg Foundation, Diabetes Research Foundation of Finnish Diabetes Association and Tampere University Hospital Supportting Foundation. M.A.-K. works in a Unit that is supported by the University of Bristol and UK Medical Research Council (MC_UU_12013/1). The Baker Institute is supported in part by the Victorian Government’s Operational Infrastructure Support Program.

## Author contributions

P.K., J.A., J.K., J.T., R.S. and J.M. designed the analyses. V.K. performed the statistical analyses. J.A., J.T., V.K., J.K., J.M., S.K.-K. and P.K. analyzed the data. M.-R.J.,J.A., S.K.-K., T.T., M.A.-K. and O.T.R. designed individual studies. J.A., J.T., R.S., E.Y.D., J.K.,T.T., M.A.-K., K.P., M.K., E.R., T.L., O.T.R., M.-R.J. and P.S. collected and generated data in individual studies. P.K., M.-R.J., J.A., J.T., V.K., J.K., J.M. and S.K.-K. wrote the manuscript.

## Competing financial interests

The authors declare no competing financial interests.

## Materials and Correspondence

Correspondence should be addressed to P.K. or M.-R.J. and request for materials to M.-R.J.

## Methods

### Study populations

We tested our hypothesis in two Finnish populations, Northern Finland Birth Cohort 1966 (NFBC1966)^1, 2^ and Cardiovascular Risk in Young Finns Study (YFS)^3^. The NFBC1966 originally included 12,068 mothers who gave birth to 12,231 live born children (96.3% of all estimated births in the two northernmost provinces of Finland, Oulu and Lapland, during the year 1966). The data collection started in 1965 when the mothers were pregnant and so far, data has been collected at ages 1, 14, 31 and 46 years. At 31- and 46-years, the data collection included postal questionnaires and clinical examinations with anthropometric and blood pressure measurements and blood sampling (n= 3624 at 31-years and n = 5351 at 46-years, Extended data Fig. 1). Maximal exercise test was performed for a subpopulation at 31-years (n = 123).

### Blood analyses

Blood samples of NFBC1966 were taken after an overnight fasting period, centrifuged immediately and before analysing samples of 31-years were stored firstly at −20?°C and later at −80?°C, whereas samples of 46-years were analysed immediately without storing. NFBC1966 blood samples were analysed in NordLab Oulu (former name Oulu University Hospital, Laboratory), a testing laboratory (T113) accredited by Finnish Accreditation Service (FINAS) (EN ISO 15189). For YFS blood analyses, see^3^.

#### Blood haemoglobin (Hb) level and red blood cell parameters

Blood Hb and red cell parameters of NFBC1966 were determined using a Coulter STKR analyser (Beckman Coulter, Fullerton, CA, USA) at age 31-years and a Sysmex XE-2100 analyser (Sysmex Corporation, Kobe, Japan) at age 46-year. The whole blood Hb levels of NFBC1966 were determined using spectrophotometric methods, red blood cells (RBC) were measured using electric resistance detecting methods (impedance technology) with hydrodynamic focusing. At age 31-years hematocrit (HCT) was measured by the sum of RBC counted in a speci?ed volume of diluted blood and at age 46-years HCT were determined applying the RBC pulse-height detection.

#### Serum lipids

Serum cholesterol (total, HDL and LDL) and triglycerides of NFBC1966 were determined using a Hitachi 911 automatic analyser and commercial reagents (Roche, Boehringer Mannheim, Germany) at 31-years^4, 5^ and an enzymatic assay method (Advia 1800; Siemens Healthcare Diagnostics Inc., Tarrytown, NY, USA) at 46-years.

#### High sensitivity C-reactive protein

High sensitivity C-reactive protein (CRP) in NFBC1966 was analysed by an immunoenzymometric assay (Medix Biochemica, Espoo, Finland) at 31-years and by an immune nefelometric assay at 46-years (BN ProSpec, Siemens Healthcare Diagnostics Inc., Newark, DE, USA).

#### Glucose and insulin levels and fasting indices

At age 31-years fasting plasma glucose and serum insulin levels of NFBC1966 were determined by glucose dehydrogenase method (Granutest 250, Diagnostica Merck, Darmstadt, Germany) and by radioimmunoassay (Pharmacia Diagnostics, Uppsala, Sweden), respectively. At 46-years fasting plasma glucose and serum insulin levels of NFBC1966 were analysed by an enzymatic dehydrogenase method (Advia 1800, Siemens Healthcare Diagnostics, Tarrytown, NY, USA) and by a chemiluminometric immunoassay (Advia Centaur XP, Siemens Healthcare Diagnostics, Tarrytown, NY, USA), respectively.

To evaluate the insulin resistance and β-cell function we calculated fasting indices HOMA-IR (fasting plasma glucose × fasting serum insulin / 22.5) and HOMA2-β ((20 × fasting serum insulin) / (fasting plasma glucose – 3.5) × 100).

#### Oral glucose tolerance test and diabetes

A two hour oral glucose tolerance test (OGTT) was performed after overnight (12 hour) fasting period for NFBC1966 at 46-years (n = 4446). Medication for diabetes or just before test measured capillary blood glucose level >8.0 mmol/l were used as exclusion criteria. Both serum insulin and plasma glucose were measured at baseline and 30, 60 and 120 minutes after 75g glucose intake. OGTT glucose and insulin values were used to calculate insulin and glucose area under curve (glucose-AUC and insulin-AUC), Matsuda index for insulin sensitivity (10 000 × ((fasting plasma glucose × fasting serum insulin) × ((fasting plasma glucose + 30 min plasma glucose + 60 min plasma glucose + 120 min plasma glucose) / 4) × ((fasting serum insulin + 30 min serum insulin + 60 min serum insulin + 120 min serum insulin) / 4)))^6^.

#### Serum metabolomics

Metabolic measures were obtained using the same high-throughput serum NMR metabolomics platform in the same analysis laboratory as described previously^7^. This methodology provided information on 228 serum measures, including lipoprotein subclass distribution and lipoprotein particle concentration, low molecular weight metabolites, such as amino acids, 3-hydroxybutyrate and creatinine, and detailed molecular information on serum lipids. The metabolite data provided by this platform has been utilized in various studies in epidemiology and genetics that have been recently reviewed^8^. The method provides metabolic measures in concentration units facilitating interpretation and replication of findings.

### Anthropometric measurements

Body weight and height and waist and hip circumferences of NFBC1966 participants were measured at 31- and 46-years. Body weight was measured with digital scale, which was calibrated regularly. Height was measured twice (mean of the two measurements was used) by using standard and calibrated stadiometer. Finally, body mass index (BMI) was calculated as the ratio of weight and height squared. Waist and hip circumferences were measured twice and mean of the two measurements was used. The waist-hip ratio was assessed as the ratio between circumferences of the waist (at the level midway between lowest rib margin and the iliac crest) and the hip (at the widest trochanters). In addition, body fat mass, fat percentage, muscle mass and visceral fat area of NFBC1966 participants at 46-years were measured by InBody 720 bioelectrical impedance analyser (Biospace Co., Ltd., Seoul, Korea). All anthropometric measurements were done after overnight (12h) fasting period. For YFS, see^3^.

### Brachial blood pressure measurement

In NFBC1966 brachial systolic and diastolic blood pressure was measured two times at 31-years and three times at 46-years with 1 min interval after 15?min of rest on the right arm of the seated participants using an automated oscillometric blood pressure device and appropriately sized cuff (Mercury sphygmomanometer at 31-years and Omron Digital Automatic Blood Pressure Monitor Model M10-IT at 46-years). Finally, the mean of two lowest systolic values and their diastolic values was used in the analyses.

### Confounders and exclusions

Smoking and physical activity were used as confounders. NFBC1966 participants reported their smoking habits at 31- and 46-years and participants we categorised them as never, former and current smokers at both time points. Physical activity was inquired and categorised as inactive, moderate or high. Individuals using combined oral contraceptives, hormone replacement therapy or serum lipids lowering drugs were omitted from the metabolomics analyses, as these have been shown to greatly influence these measures (total n = 957). Individuals with known diabetes before clinical examinations were omitted from analyses. Diabetes was defined according to self-reported diagnoses and medications, hospital outpatient and inpatient registers and medication registers from Social Insurance Institution of Finland. For YFS, see^3^.

### Analyses in mice

All animal experiments were performed according to protocols approved by the Provincial State Office of Southern Finland (license number ESAVI/6154/04.10.07/2014). The investigations conform to the Guiding Principles for research involving animals. Male C57Bl/6N mice (Charles River, Germany) were maintained in plastic cages in a constant 21°C temperature with a 12 h light-dark cycle and had free access to food. After weaning, the mice were fed Teklad Global 18% Protein Rodent Diet (18.6% protein, 6.2% fat, 44.2% carbohydrates; Envigo) and at 5-weeks of age the mice were transferred to Teklad Global 19% Protein Extruded Rodent Diet (19.0% protein, 9.0% fat, 44.9% carbohydrates; Envigo) to slightly support the increase in adipocity upon aging. At 11-months of age the mice were subjected to glucose tolerance test (GTT). GTT was performed on mice fasted for 12 h and anesthetized with fentanyl/fluanisone and midazolam. Mice were injected intraperitoneally with 1 mg/g glucose, and blood glucose concentrations were monitored with a glucometer. Serum insulin values were determined with Rat/Mouse Insulin ELISA kit (EZRMI-13K; Millipore), and HOMA-IR scores were calculated from the glucose and insulin values. The mice were weighed, sampled for hemoglobin determined (by HemoCue Hb 201 hemoglobin meter) and sacrificed at the age of one year. For the venesection experiment 3-month-old C57Bl/6N male mice fed Teklad Global 18% Protein Rodent Diet were fasted for 12 h and anesthetized as above. Mice were weighted and their fasting blood glucose values and Hb levels were determined. Samples were collected for determination of serum fasting insulin, total cholesterol and triglyceride levels. Mice were then injected intraperitoneally with 2 mg/g glucose and blood glucose concentration after 2 h was monitored. The mice were then subjected to venesection of 0.2 ml. After 14 days the same analyses (body weight, Hb, fasting glucose, fasting insulin, 2-h glucose in GTT, total cholesterol and triglycerides) were repeated and the mice were sacrificed. Statistical analyses for comparison between two groups were done with two-tailed Student’s *t test*. Pearson’s correlation coefficient was calculated to compare linear dependences between two variables. Areas under the curve were calculated by the summary measures method.

### Statistical analyses

Regression models were used to assess the associations between Hb (main explanatory variable) and anthropometric and metabolic variables (outcome). To adjust for potential confounding factors, smoking, physical activity and sex were included as explanatory variables in the regression models. For the analyses using YFS data, age was also included as an explanatory variable. The analyses were conducted including or excluding observations with an absolute value more than 3 standard deviations off from the mean of Hb or the (potentially transformed) outcome variable. Linear associations were assumed for all continuous variables. To ease the comparison of magnitudes of association across outcomes, the effect size estimates are reported in standard deviation units. To account for multiple comparisons in the anthropometric and metabolome-wide analyses, principal component analysis was used to estimate the number of independent tests for a Bonferroni-style multiple testing correction. As 23 principal components explained 95% of the variation in the anthropometric and metabolic outcomes, a *P* value of 0.002 (0.05/23) was used as a nominal threshold for significance in these analyses. R 3.3.2 software was used for all statistical analyses (R Core Team (2013): R: a language and environment for statistical computing, Vienna, Austria, http://www.R-project.org/).

#### Anthropometric measures

BMI was log-transformed to eliminate skewness and heteroscedasticity in model residuals. Height was included as an additional explanatory variable in the regression models, excluding the model for BMI. The analyses were first conducted separately for the 46-age year follow-up in NFBC1966 and for YFS, and then meta-analyzed for those outcomes available in both cohorts using an inverse-variance weighted random-effects model. The meta-analysis effect sizes were calculated and reported in the units in which they were measured. For the NFBC1966 dataset, similar regression models were conducted for red blood cell counts (Eryt) and hematocrit (Hct) instead of Hb, to examine the association between these measures and anthropometric variables.

#### Metabolic parameters

Fasting insulin levels, Matsuda and HOMA indices, AUCs and trigyceride and hsCRP levels were log-transformed to ensure symmetry and homoscedasticity of model residuals. In the hsCRP variable, including zeros in the values, the transformation used was *log*(*x* + *c*/2), where *c* = *min*(*x*_*x*≠0)_, *i*.*e*. the smallest non-zero value in the variable. The analyses were conducted for the 46-year follow-up data in NFBC1966 with both including and excluding BMI as an additional explanatory variable.

#### NMR metabolomics

In total, 150 metabolites from the NMR metabolomics panel were analyzed (Extended data Table 8). Of these, 144 were measured in concentrations, and cube-root transformation was applied to these variables^9^. The rest of the variables were left untransformed, except the degree of unsaturation, which had a notably skewed distribution and thus log-transformed to ensure homoscedasticity in model residuals. The users of combined oral contraceptives, hormone replacement therapy and lipid lowering drugs (total n = 957) were omitted because their influence on these measures^10,11^. The analyses were conducted first separately in the NFBC1966 (at 46 years) and YFS datasets, and then meta-analyzed using inverse-variance weighted random-effects models. BMI was included as an additional explanatory variable.

#### Longitudinal models in NFBC1966

Extended linear models using generalized least squares were used to examine the longitudinal change in association between Hb levels and the outcomes available at two time points in the NFBC1966. Time of measurement (ages of 31 years or 46 years) was included in the model as a categorical variable, and interaction terms with time and all other variables (excluding sex and height where applicable) were included in the model to allow varying effects between cross-sectional measurements. The error terms were allowed to be correlated within each individual and heteroskedastic between the time points. These analyses were conducted using the *gls()* function available in the *nlme* package^12^. Due to the different methods used to measure hsCRP at 31 and 46 years, an inverse normal rank transformation was applied to set the values on a comparable scale. As the glucose concentrations at the NFBC1966 31-year follow-up were measured from whole blood these values were transformed to corresponding concentrations in plasma^13^ for compatibility with the data at 46-year follow-up.

#### Cross-sectional and longitudinal analysis of Hb levels on metabolites

To analyse both cross-sectional and longitudinal effects of Hb levels on the metabolites, Hb levels were decomposed to mean Hb 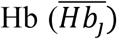 for each individual *j* and variation of Hb from the individual mean 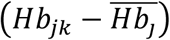 for each individual *j* at time point *k* (1 = 31 years, 2 = 46 years). The following model was fit for each metabolite separately:

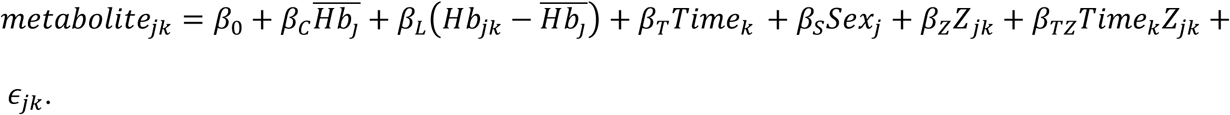

This model allows simultaneous estimation of the cross-sectional (between-subject) effect *β*_*c*_ and the longitudinal (within-subject) effect *β*_*L*_^14^, while controlling for potentially time-varying covariates *Z*_*jk*_, in this case BMI, smoking and physical activity. The error terms ϵ_*jk*_ were allowed to be correlated within each individual and heteroskedastic between the time points. These analyses were conducted using the *gls()* function available in the *nlme* package^2^.

#### Maximal exercise test subsample in NFBC1966

Regression models of the effects of Hb levels on oxygen consumption at rest divided by height (VO2rest/h), VO_2_rest/h on body fat percentage and Hb levels on body fat percentage were conducted for the subset of NFBC1966 participants at 31 years who underwent a maximal exercise test with direct measurement of oxygen consumption (M901, ergospirometer, Medikro, Finland)^15^. These analyses included sex as an additional explanatory variable.

#### Mendelian randomization

Bidirectional Mendelian randomization (MR) was conducted to examine a potential causality of Hb levels on BMI and *vice versa*. In the MR approach, genetic variants predicting an observational exposure are used as proxies for the exposure. If these variants fulfil instrumental variable assumptions^16^, then association between the variants and the outcome can give evidence for causation, eliminating problems of confounding or reverse causation. Using a *P* value threshold of 1 × 10^-8^, 56 and 17 single nucleotide polymorphisms (SNPs) from independent genetic loci associated with BMI and Hb, respectively, were identified from European ancestry genome-wide meta-analyses for the two traits^17,18^. All BMI- and Hb-associated SNPs were available in the Hb and BMI meta-analysis results, respectively. After harmonizing the effect alleles for each SNP, the effect estimates and their standard errors of BMI-related SNPs were extracted from the summary statistics available for Hb and *vice versa*. Causal effects were estimated from these summary data using both the inverse-variance weighted (IVW) method^19^ and MR-Egger regression^20^. The IVW method is a weighted regression of SNP-outcome on SNP-exposure coefficients, weighted by the inverse of the squared SNP-outcome standard errors. The slope of this model is an estimate for the causal effect. The intercept is constrained at zero, implying the assumption that the genetic data predicts the outcome only through the exposure. MR-Egger regression relaxes this assumption by including an intercept term, which is an estimate for the average pleiotropic effect of a SNP on the outcome. Testing whether this intercept equals zero serves as a test for directional pleiotropy. Assuming that magnitude of the SNP-exposure associations across all variants are independent of their pleiotropic effects, the slope in MR Egger regression provides a consistent estimate of the causal effect. Sources of bias to these causal effect estimates include deviations of the SNP-exposure estimates from the true associations, use of weak instruments, and participant overlap in the meta-analysis from where the effect estimates were calculated. In order to quantify the potential bias, relative bias estimates (the reciprocals of the F parameters^21^) were calculated for the IVW method estimates and 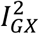 statistics^22^ for the MR Egger estimates.

#### Polygenic risk score analyses

We conducted polygenic risk score (PRS) analyses to further examine the association between Hb, BMI and lipids. We generated PRSs for all exposure-outcome combinations using exposure GWAS results^23-26^ for SNP weights, optimised for each outcome in NFBC1966 at 46 years. We used PRSice software^27^ for PRS calculation, using a clumping distance of 250kb and an *r*^*2*^ threshold of 0.1. Ten first genetic principal components were added as covariates to account for population stratification. PRS was calculated for a range of p-value cut-offs (from 5×10^-8^ to 1 by 0.05 increments) and we report the PRS using a p-value cut-off giving the highest *R*^*2*^ with the target outcome. As NFBC1966 is part of all base GWASs used for PRS calculation, we might expect some inflation in the PRS variance explained. As a sensitivity test, we tested optimising PRS using BMI GWAS results from UKBiobank (http://www.nealelab.is/blog/2017/9/11/details-and-considerations-of-the-uk-biobank-gwas) on Hb in NFBC1966 at 46 years. The *R*^*2*^ for the PRS using the external GWAS results was 0.35 % (positive association, p = 4e-4), compared to 0.17 % (positive association, p = 0.01) with GWAS results with NFBC1966 included, showing no evidence for inflation in the variance explained.

#### Gene Set Enrichment Analysis

To examine the hypothesized activation of hypoxia response pathway, a Gene Set Enrichment Analysis (GSEA)^28^ was conducted for the subset of YFS participants at 34-49 years with Hb levels and whole-blood genome-wide expression profiling data (n = 1636)^29^. The tested hypoxia response pathway included 22 genes induced at least four-fold by hypoxia in human monocytes (Extended data Table 11)^30^. The GSEA was performed between YFS participants in lowest (Hb <132, n = 392) and highest (Hb > 152, n = 371) quartile of Hb distribution. The analysis was done adjusted for age, sex, BMI, YFS study center, leukocyte and thrombocyte counts, five first principal components of transcriptomics data and three technical variables from microarray hybridizations (chip, plate and well). The analysis was done with residual gene expression profiles calculated by linear regression of each gene expression probe and above mentioned variables. A nominal *P* value < 0.05 was considered significant.

## Auvinen et al. Lower hemoglobin levels associate with lower body mass index and healthier metabolic profile Extended data

**Extended data Table 1.**
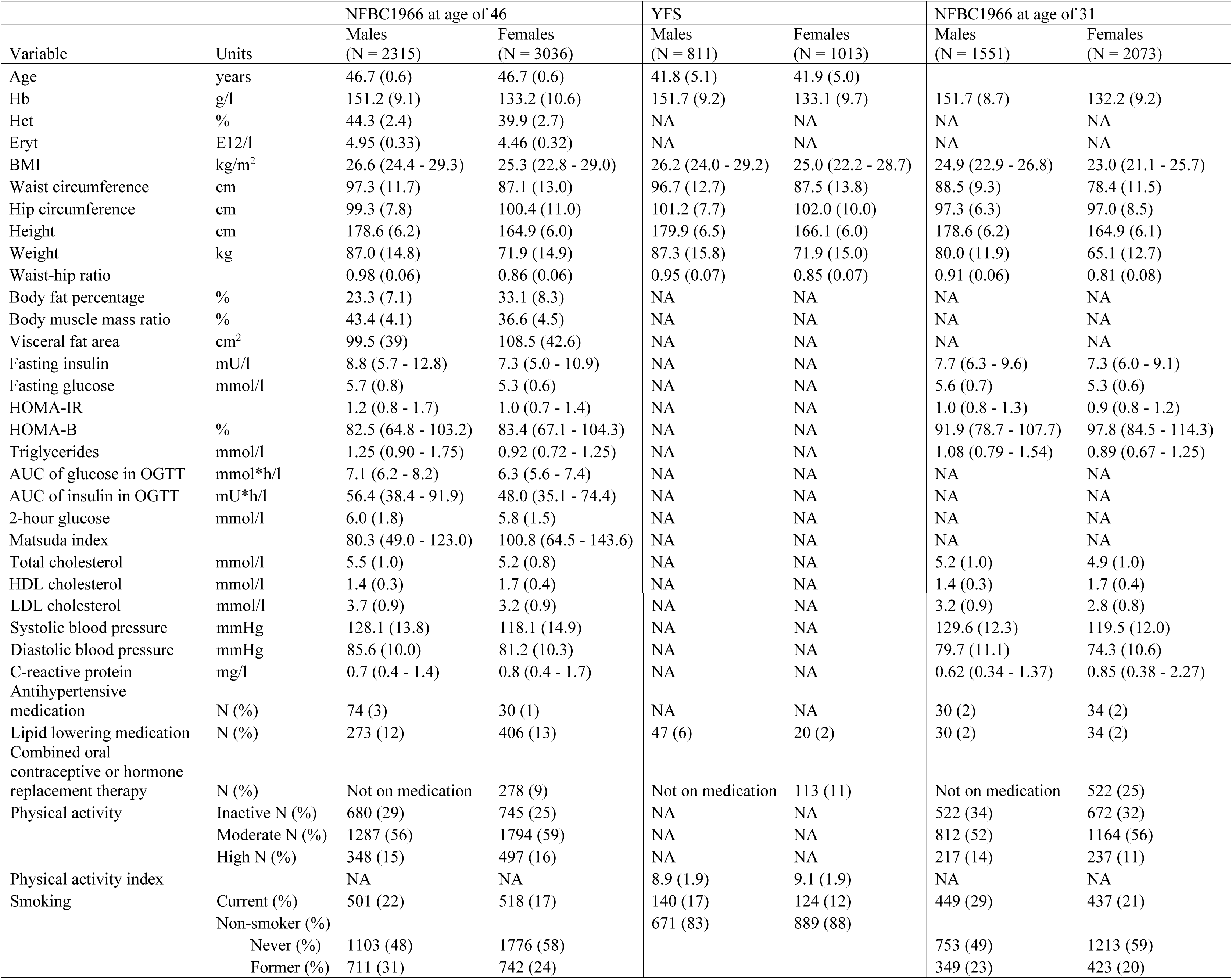
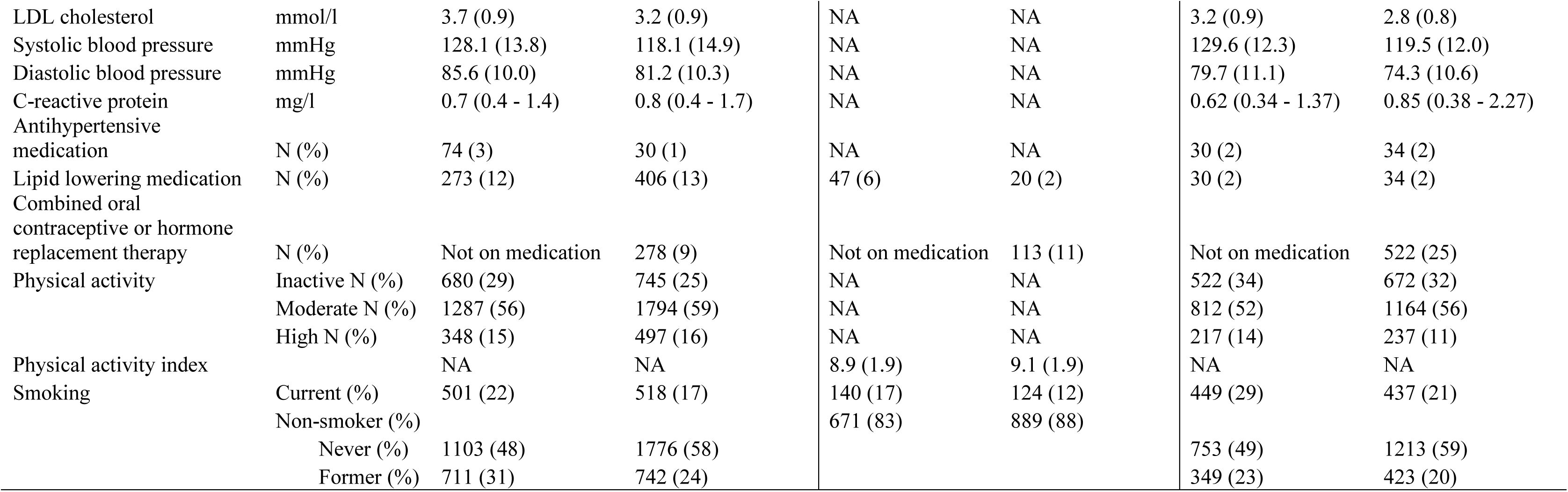
Characteristics of the NFBC1966 and YFS study populations. The numbers indicate mean and those in parentheses SD or 95% CI, if not indicated otherwise. NA, not available.

**Extended data Table 2.**
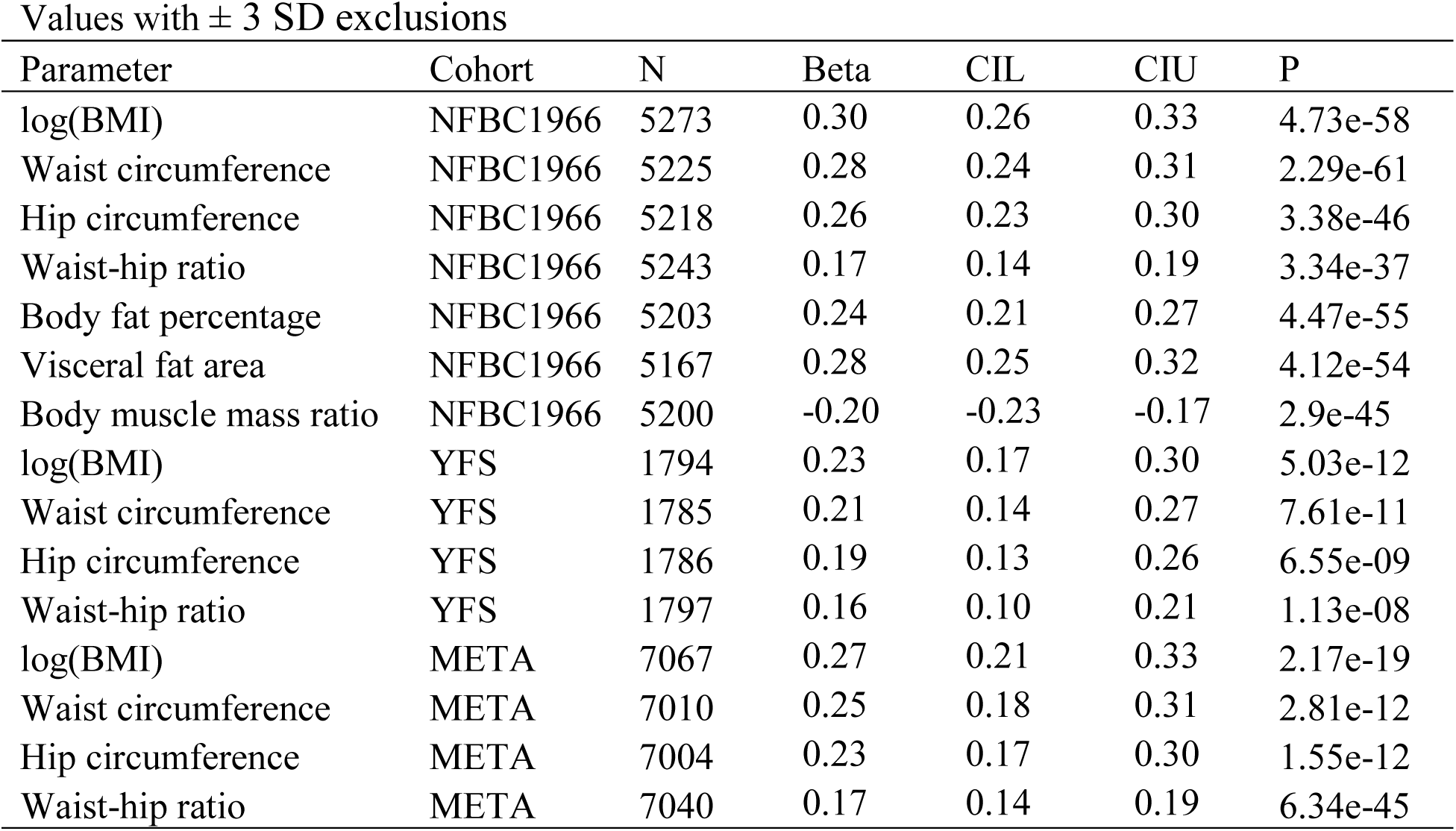
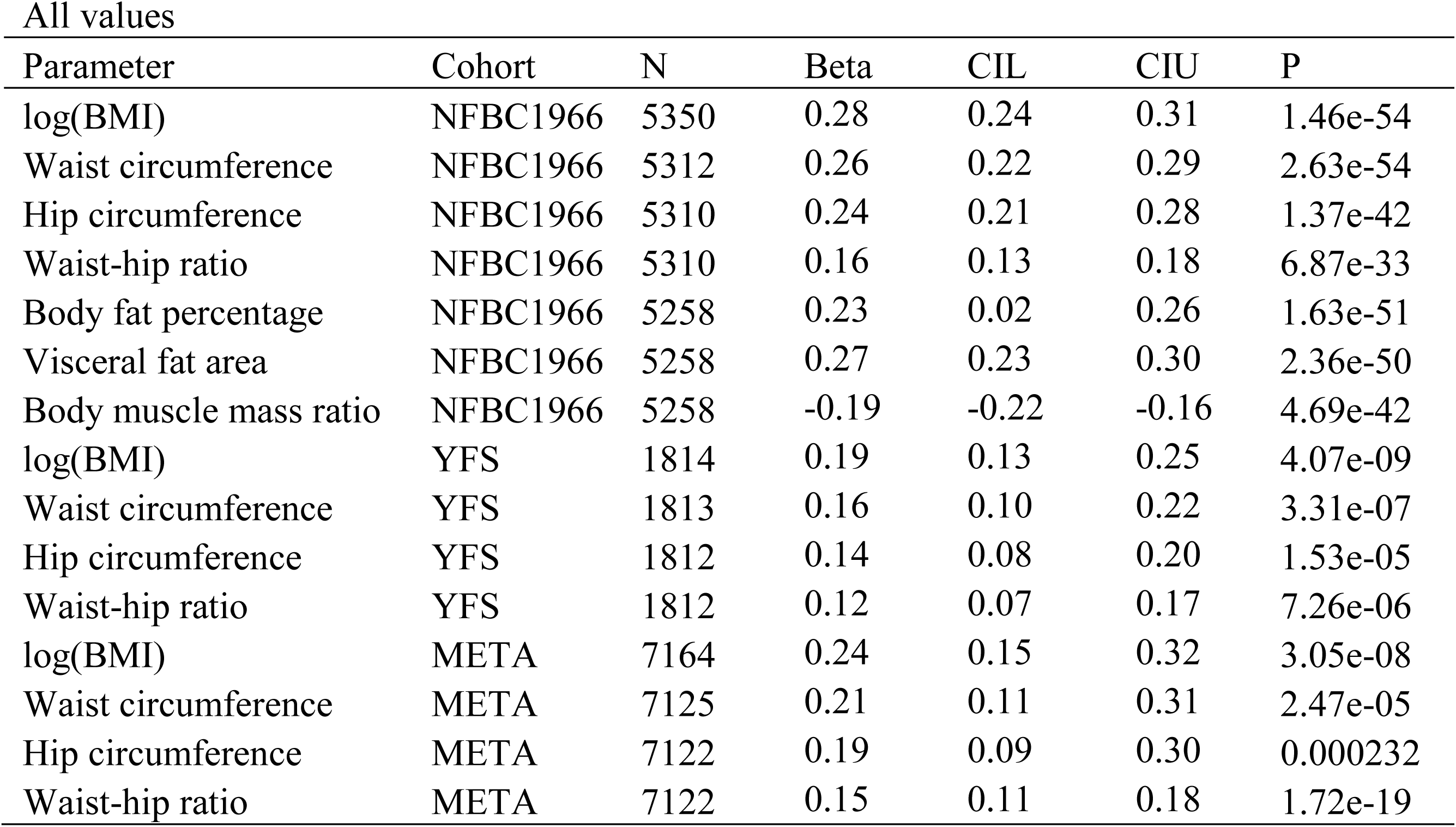
Effect sizes for association of Hb levels with BMI and other anthropometric measures. Number of participants (N) in the statistical analyses, effect sizes (Beta) in units of 1-SD change in anthropometric measures by 1-SD change in Hb, lower and upper 95% confidence intervals (CIL, CIU) and *P* values of the study population in analyses. For other parameters than BMI the BMI adjusted effects sizes are shown. META indicates meta-analysis of NFBC1966 at age of 46 and YFS.

**Extended data Table 3.**
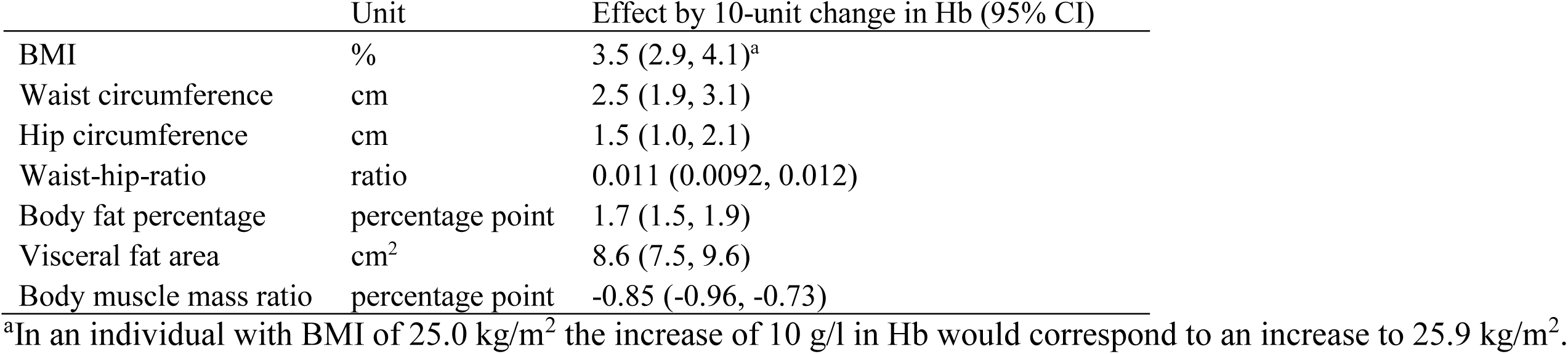
Examples of effects in Hb level changes on BMI and other anthropometric measures. The data are from the meta-analysis presented in Extended data Table 2 (values with ± 3 SD exclusions).

**Extended data Table 4.**
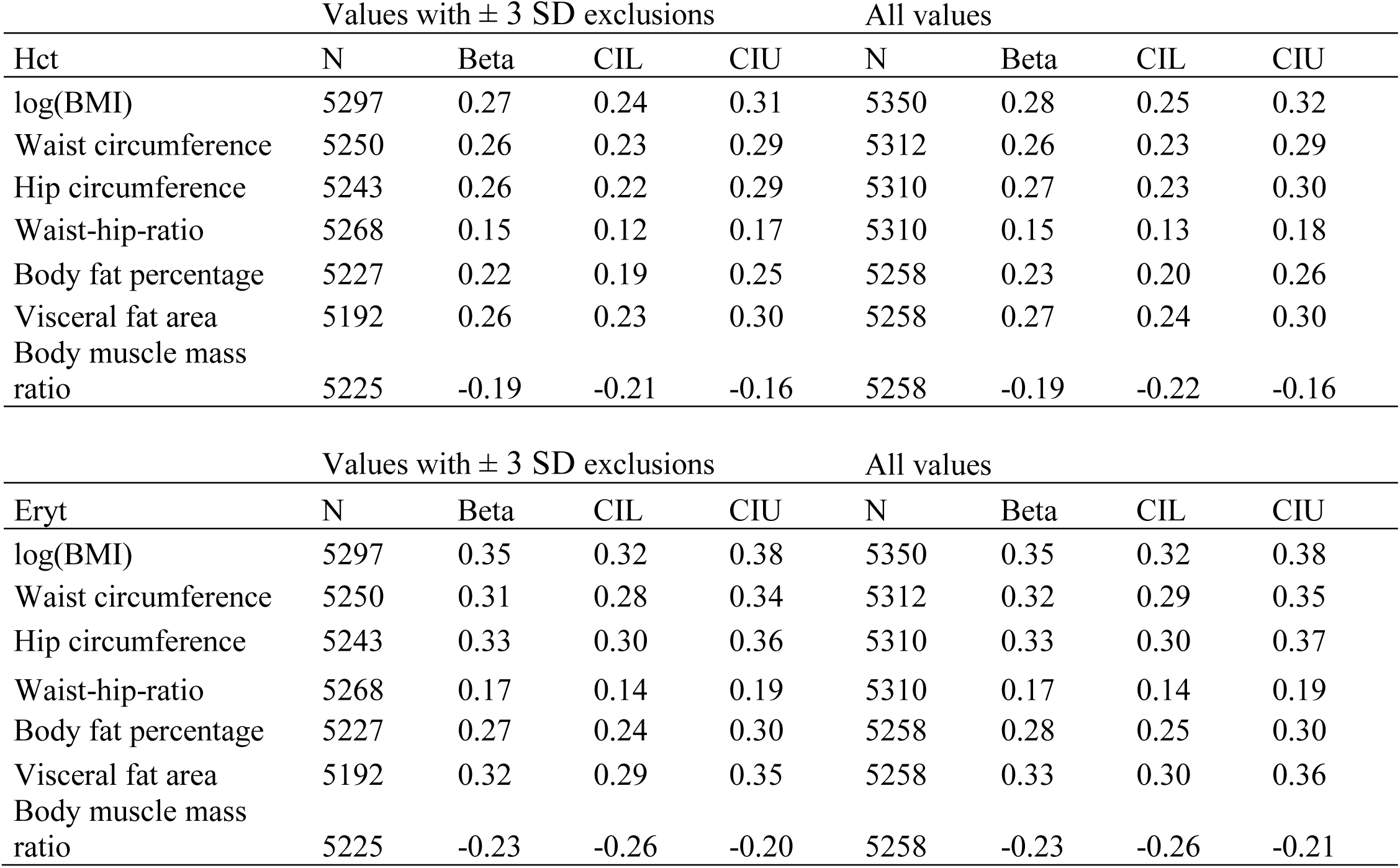
Effect sizes for association of hematocrit levels and red blood cell counts with BMI and other anthropometric measures. Number of participants in the statistical analyses (N), effect sizes (Beta) in units of 1-SD change in anthropometric measures by 1-SD change in hematocrit (Hct) or red blood cell counts (Eryt) and lower and upper 95% confidence intervals (CIL, CIU) of the NFBC1966 at age of 46.

**Extended data Table 5.**
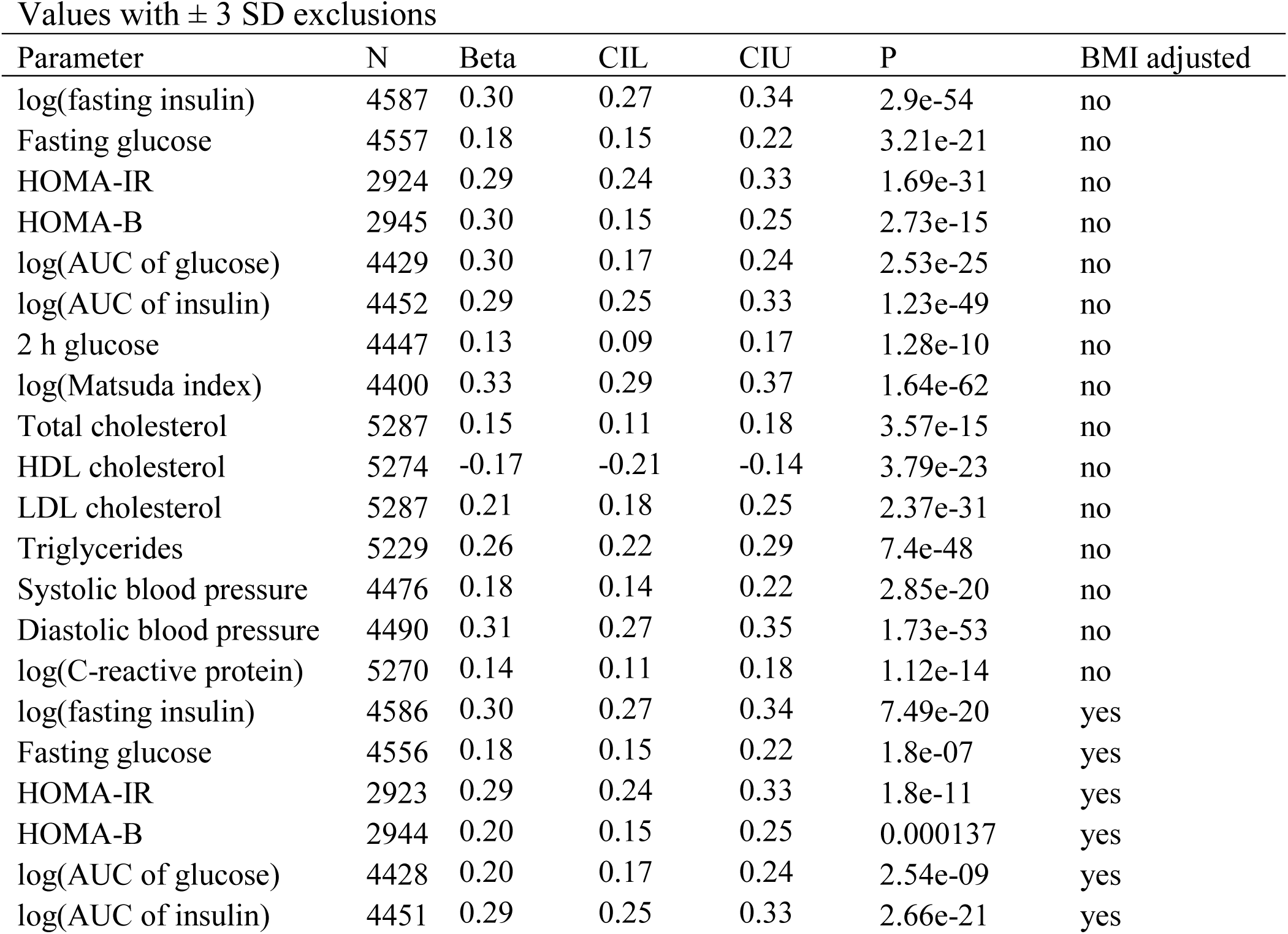
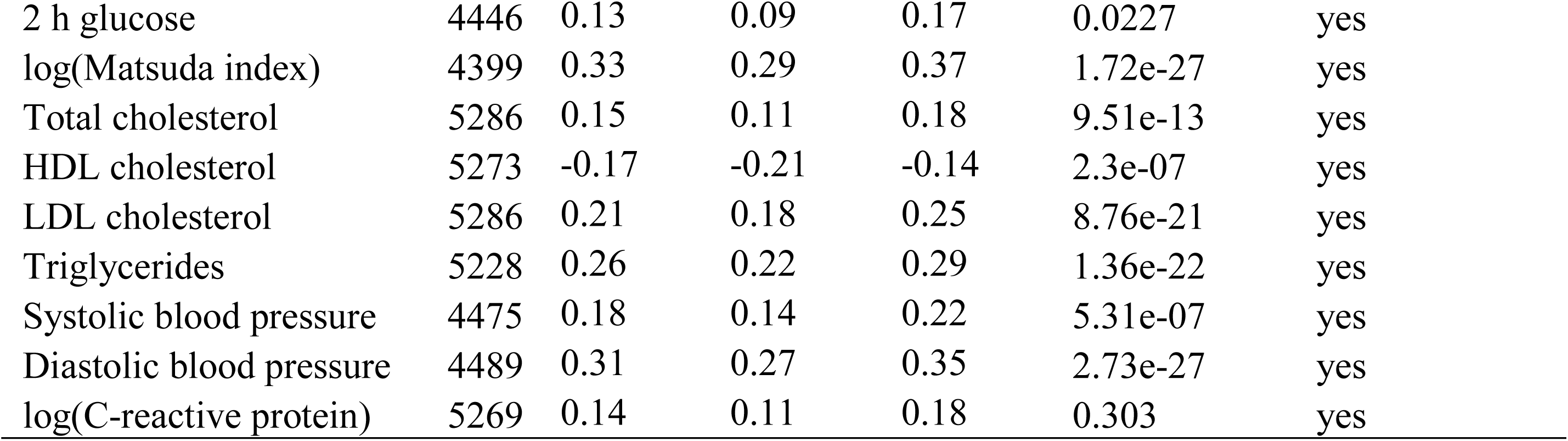
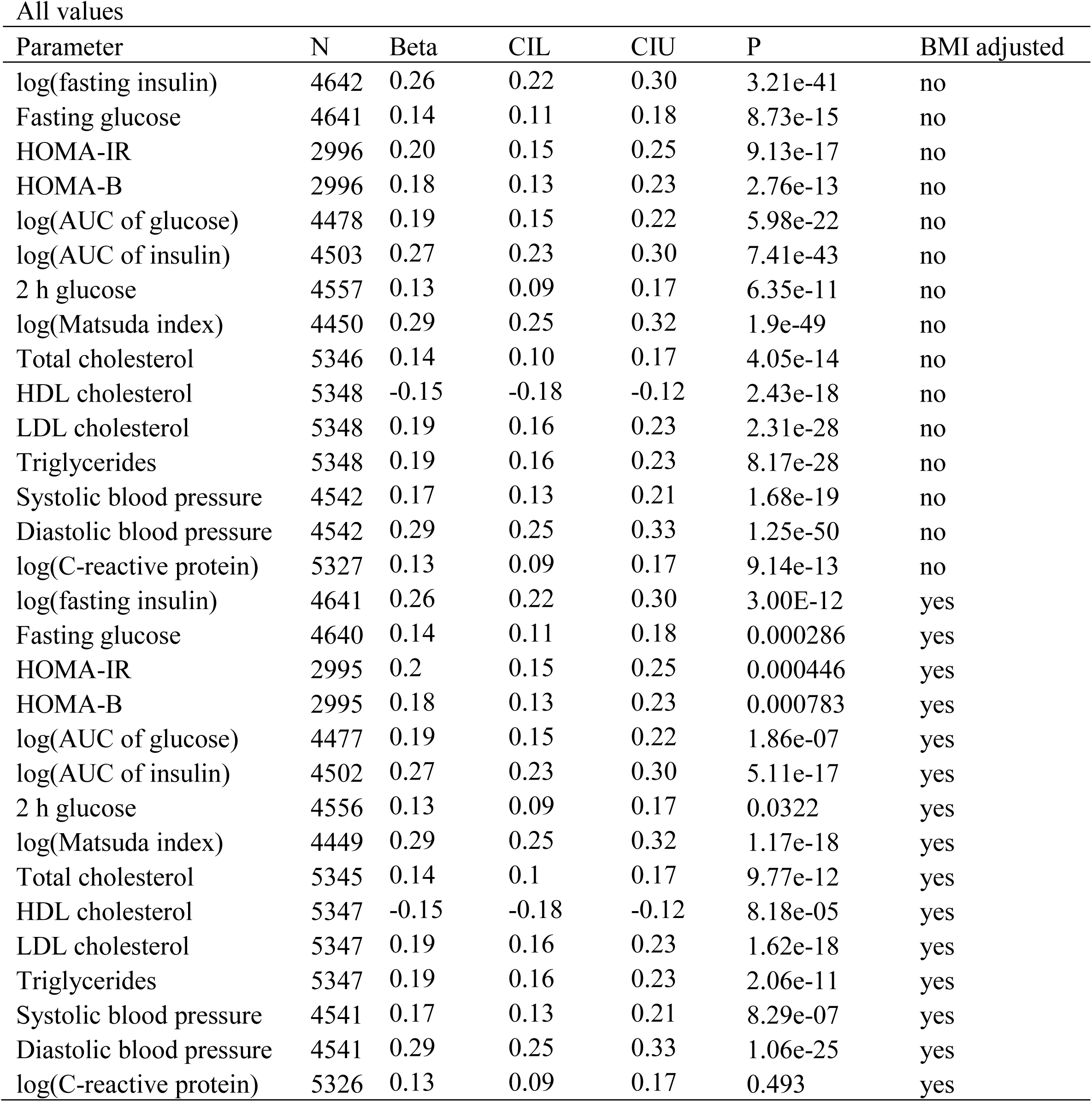
Effect sizes for association of Hb levels with glucose and lipid metabolism, blood pressure and inflammatory parameters. Number of participants (N) in the statistical analyses, effect sizes (Beta) in units of 1-SD change in the indicated parameter by 1-SD change in Hb, lower and upper 95% confidence intervals (CIL, CIU) and *P* values of the NFBC1966 at age of 46. When indicated the data were adjusted for BMI. Users of antihypertensive (n = 815) drugs were removed from the systolic and diastolic blood pressure analyses.

| Values with $\pm 3$ SD exclusions | | | | | | |
| --- | --- | --- | --- | --- | --- | --- |
| Parameter | N | Beta | CIL | CIU | P | BMI adjusted |
| log(fasting insulin) | 4587 | 0.30 | 0.27 | 0.34 | 2.9e-54 | no |
| Fasting glucose | 4557 | 0.18 | 0.15 | 0.22 | 3.21e-21 | no |
| HOMA-IR | 2924 | 0.29 | 0.24 | 0.33 | 1.69e-31 | no |
| HOMA-B | 2945 | 0.30 | 0.15 | 0.25 | 2.73e-15 | no |
| log(AUC of glucose) | 4429 | 0.30 | 0.17 | 0.24 | 2.53e-25 | no |
| log(AUC of insulin) | 4452 | 0.29 | 0.25 | 0.33 | 1.23e-49 | no |
| 2 h glucose | 4447 | 0.13 | 0.09 | 0.17 | 1.28e-10 | no |
| log(Matsuda index) | 4400 | 0.33 | 0.29 | 0.37 | 1.64e-62 | no |
| Total cholesterol | 5287 | 0.15 | 0.11 | 0.18 | 3.57e-15 | no |
| HDL cholesterol | 5274 | -0.17 | -0.21 | -0.14 | 3.79e-23 | no |
| LDL cholesterol | 5287 | 0.21 | 0.18 | 0.25 | 2.37e-31 | no |
| Triglycerides | 5229 | 0.26 | 0.22 | 0.29 | 7.4e-48 | no |
| Systolic blood pressure | 4476 | 0.18 | 0.14 | 0.22 | 2.85e-20 | no |
| Diastolic blood pressure | 4490 | 0.31 | 0.27 | 0.35 | 1.73e-53 | no |
| log(C-reactive protein) | 5270 | 0.14 | 0.11 | 0.18 | 1.12e-14 | no |
| log(fasting insulin) | 4586 | 0.30 | 0.27 | 0.34 | 7.49e-20 | yes |
| Fasting glucose | 4556 | 0.18 | 0.15 | 0.22 | 1.8e-07 | yes |
| HOMA-IR | 2923 | 0.29 | 0.24 | 0.33 | 1.8e-11 | yes |
| HOMA-B | 2944 | 0.20 | 0.15 | 0.25 | 0.000137 | yes |
| log(AUC of glucose) | 4428 | 0.20 | 0.17 | 0.24 | 2.54e-09 | yes |
| log(AUC of insulin) | 4451 | 0.29 | 0.25 | 0.33 | 2.66e-21 | yes |
| 2 h glucose | 4446 | 0.13 | 0.09 | 0.17 | 0.0227 | yes |
| log(Matsuda index) | 4399 | 0.33 | 0.29 | 0.37 | 1.72e-27 | yes |
| Total cholesterol | 5286 | 0.15 | 0.11 | 0.18 | 9.51e-13 | yes |
| HDL cholesterol | 5273 | -0.17 | -0.21 | -0.14 | 2.3e-07 | yes |
| LDL cholesterol | 5286 | 0.21 | 0.18 | 0.25 | 8.76e-21 | yes |
| Triglycerides | 5228 | 0.26 | 0.22 | 0.29 | 1.36e-22 | yes |
| Systolic blood pressure | 4475 | 0.18 | 0.14 | 0.22 | 5.31e-07 | yes |
| Diastolic blood pressure | 4489 | 0.31 | 0.27 | 0.35 | 2.73e-27 | yes |
| log(C-reactive protein) | 5269 | 0.14 | 0.11 | 0.18 | 0.303 | yes |

| All values |  |  |  |  |  |  |
| --- | --- | --- | --- | --- | --- | --- |
| Parameter | N | Beta | CIL | CIU | P | BMI adjusted |
| log(fasting insulin) | 4642 | 0.26 | 0.22 | 0.30 | 3.21e-41 | no |
| Fasting glucose | 4641 | 0.14 | 0.11 | 0.18 | 8.73e-15 | no |
| HOMA-IR | 2996 | 0.20 | 0.15 | 0.25 | 9.13e-17 | no |
| HOMA-B | 2996 | 0.18 | 0.13 | 0.23 | 2.76e-13 | no |
| log(AUC of glucose) | 4478 | 0.19 | 0.15 | 0.22 | 5.98e-22 | no |
| log(AUC of insulin) | 4503 | 0.27 | 0.23 | 0.30 | 7.41e-43 | no |
| 2 h glucose | 4557 | 0.13 | 0.09 | 0.17 | 6.35e-11 | no |
| log(Matsuda index) | 4450 | 0.29 | 0.25 | 0.32 | 1.9e-49 | no |
| Total cholesterol | 5346 | 0.14 | 0.10 | 0.17 | 4.05e-14 | no |
| HDL cholesterol | 5348 | -0.15 | -0.18 | -0.12 | 2.43e-18 | no |
| LDL cholesterol | 5348 | 0.19 | 0.16 | 0.23 | 2.31e-28 | no |
| Triglycerides | 5348 | 0.19 | 0.16 | 0.23 | 8.17e-28 | no |
| Systolic blood pressure | 4542 | 0.17 | 0.13 | 0.21 | 1.68e-19 | no |
| Diastolic blood pressure | 4542 | 0.29 | 0.25 | 0.33 | 1.25e-50 | no |
| log(C-reactive protein) | 5327 | 0.13 | 0.09 | 0.17 | 9.14e-13 | no |
| log(fasting insulin) | 4641 | 0.26 | 0.22 | 0.30 | 3.00E-12 | yes |
| Fasting glucose | 4640 | 0.14 | 0.11 | 0.18 | 0.000286 | yes |
| HOMA-IR | 2995 | 0.2 | 0.15 | 0.25 | 0.000446 | yes |
| HOMA-B | 2995 | 0.18 | 0.13 | 0.23 | 0.000783 | yes |
| log(AUC of glucose) | 4477 | 0.19 | 0.15 | 0.22 | 1.86e-07 | yes |
| log(AUC of insulin) | 4502 | 0.27 | 0.23 | 0.30 | 5.11e-17 | yes |
| 2 h glucose | 4556 | 0.13 | 0.09 | 0.17 | 0.0322 | yes |
| log(Matsuda index) | 4449 | 0.29 | 0.25 | 0.32 | 1.17e-18 | yes |
| Total cholesterol | 5345 | 0.14 | 0.1 | 0.17 | 9.77e-12 | yes |
| HDL cholesterol | 5347 | -0.15 | -0.18 | -0.12 | 8.18e-05 | yes |
| LDL cholesterol | 5347 | 0.19 | 0.16 | 0.23 | 1.62e-18 | yes |
| Triglycerides | 5347 | 0.19 | 0.16 | 0.23 | 2.06e-11 | yes |
| Systolic blood pressure | 4541 | 0.17 | 0.13 | 0.21 | 8.29e-07 | yes |
| Diastolic blood pressure | 4541 | 0.29 | 0.25 | 0.33 | 1.06e-25 | yes |
| log(C-reactive protein) | 5326 | 0.13 | 0.09 | 0.17 | 0.493 | yes |

**Extended data Table 6.**
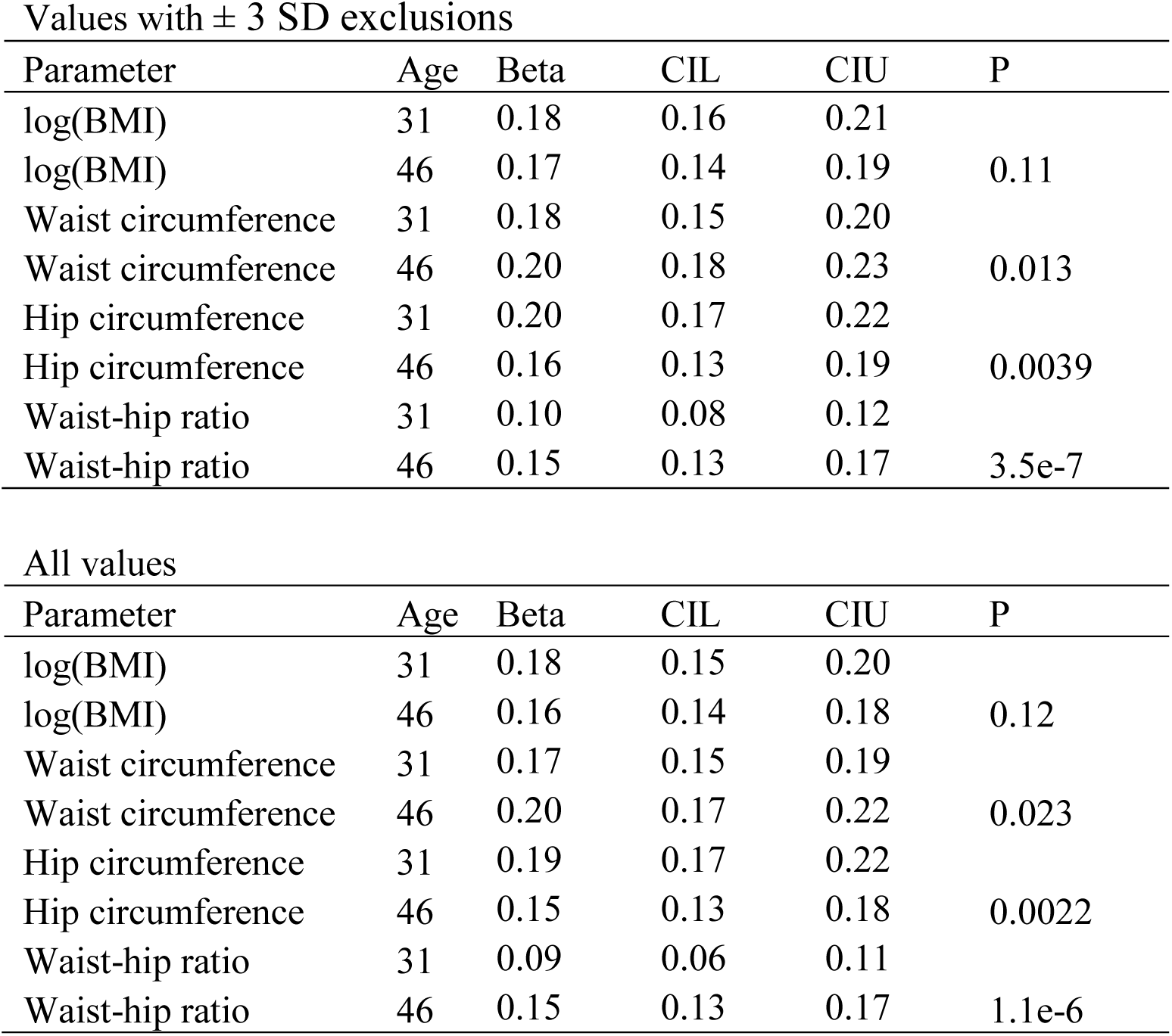
Comparison of effect sizes for association of Hb levels with BMI and other anthropometric measures at age 31 and 46 in NFBC1966. Age, effect sizes (Beta) in units of 1-SD change in the indicated parameter by 1-SD change in Hb, lower and upper 95% confidence intervals (CIL, CIU) and *P* values for the change in effect size from age 31 to 46 of the NFBC1966.

**Extended data Table 7.**
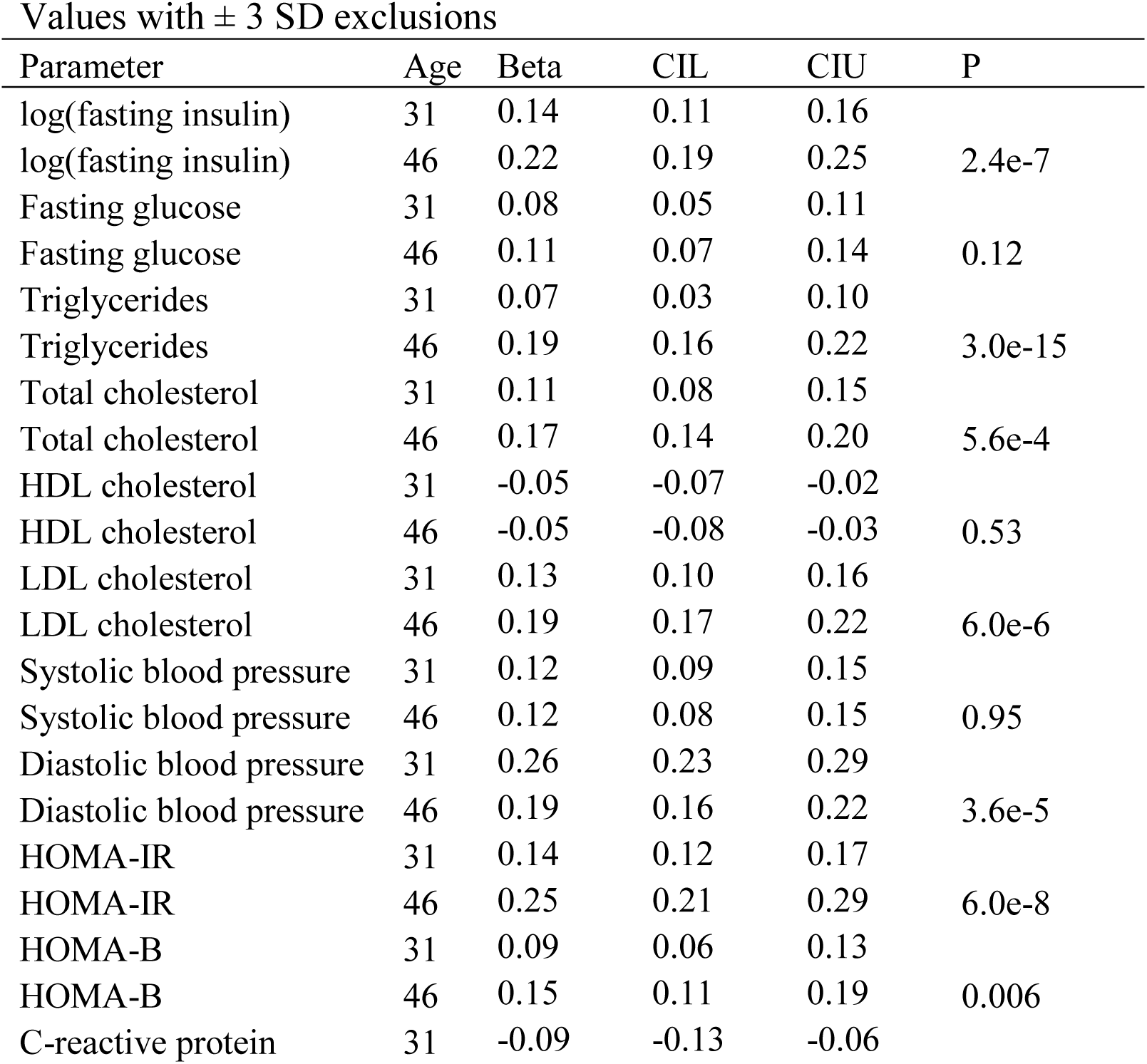
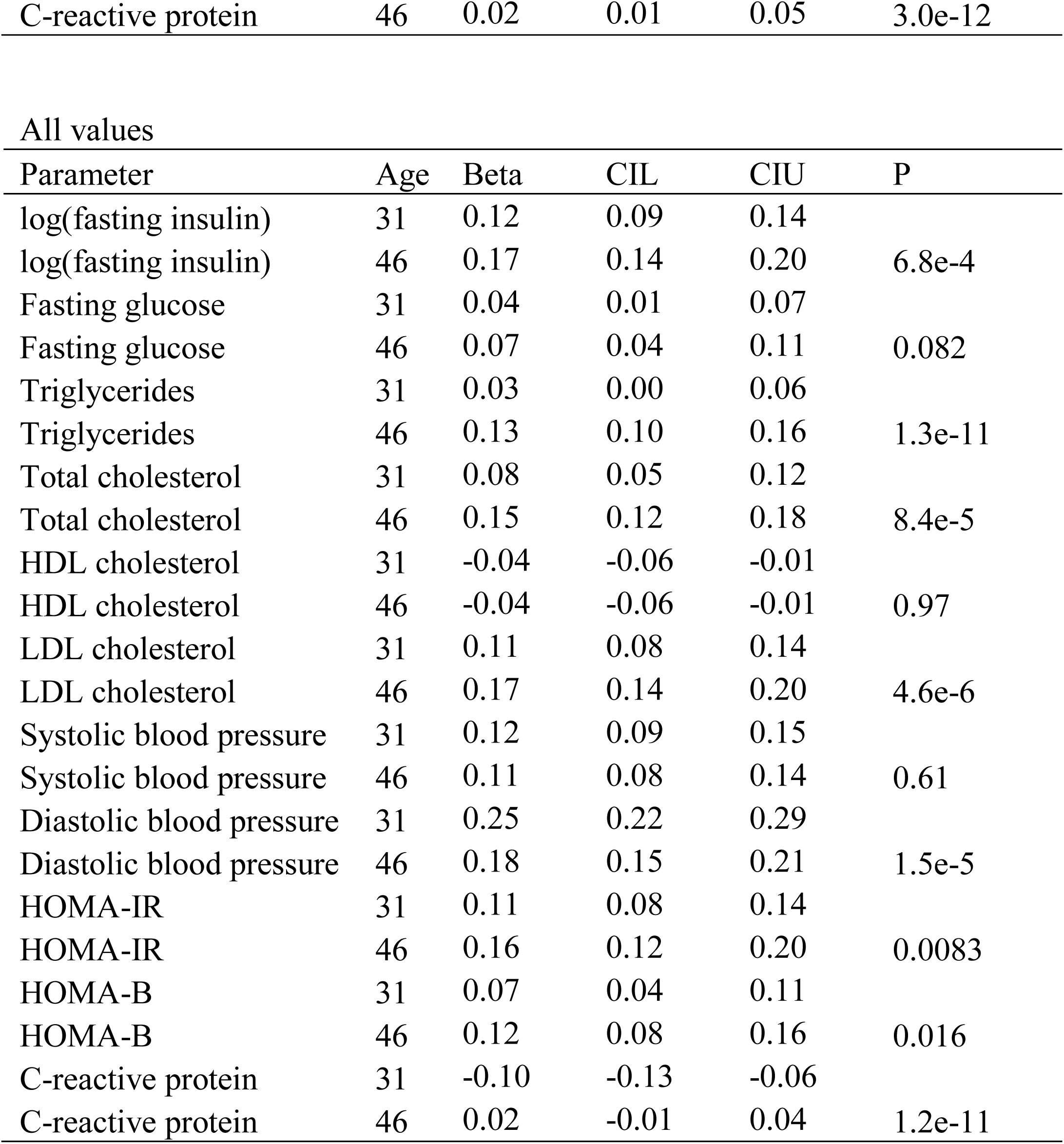
Comparison of effect sizes for association of Hb levels with glucose and lipid metabolism, blood pressure and inflammatory parameters at age 31 and 46 in NFBC1966. Age, effect sizes (Beta) in units of 1-SD change in the indicated parameter by 1-SD change in Hb, lower and upper 95% confidence intervals (CIL, CIU) and *P* values for the change in effect size from age 31 to 46 of the NFBC1966. BMI adjusted effects sizes are shown.

**Extended data Table 8.**
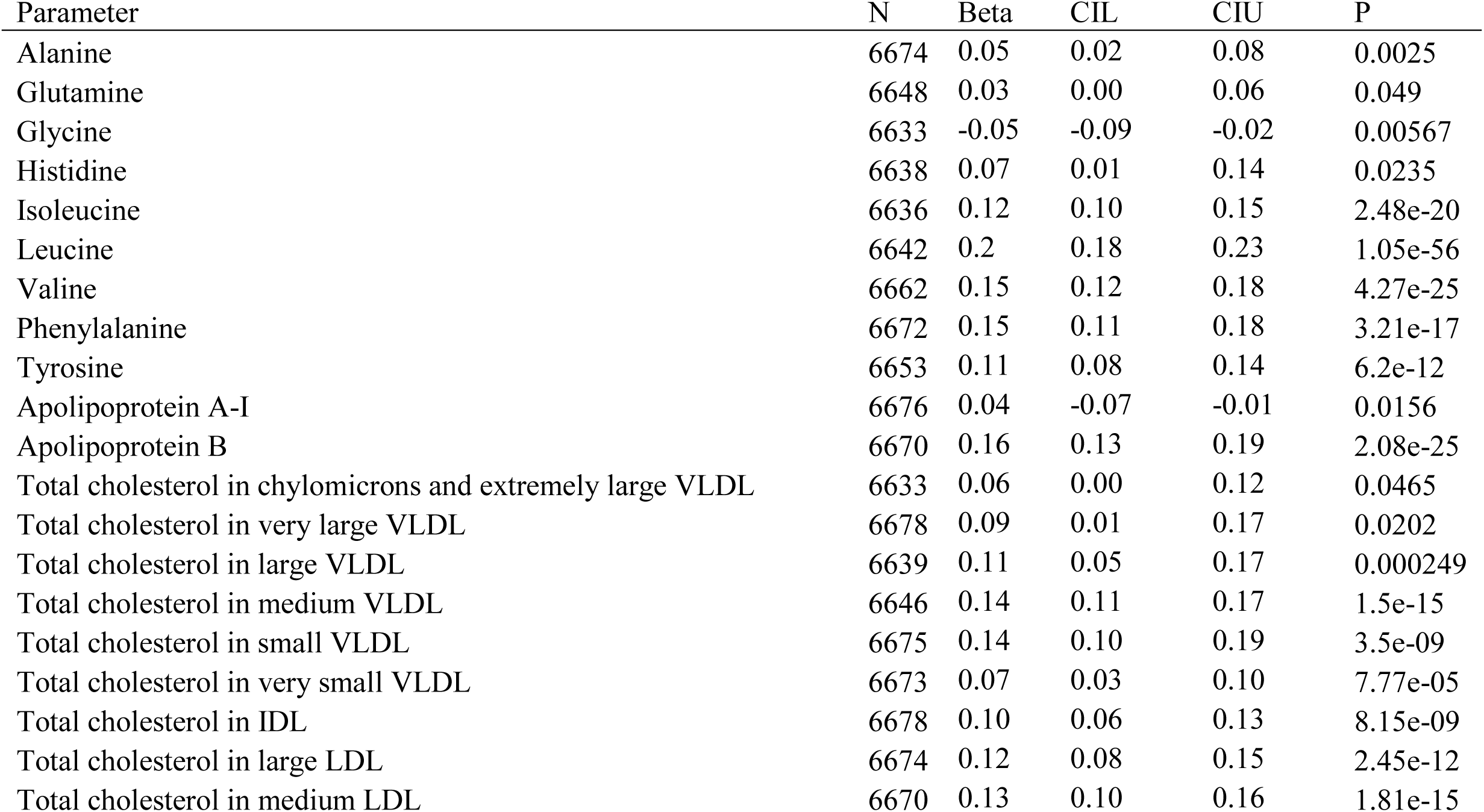
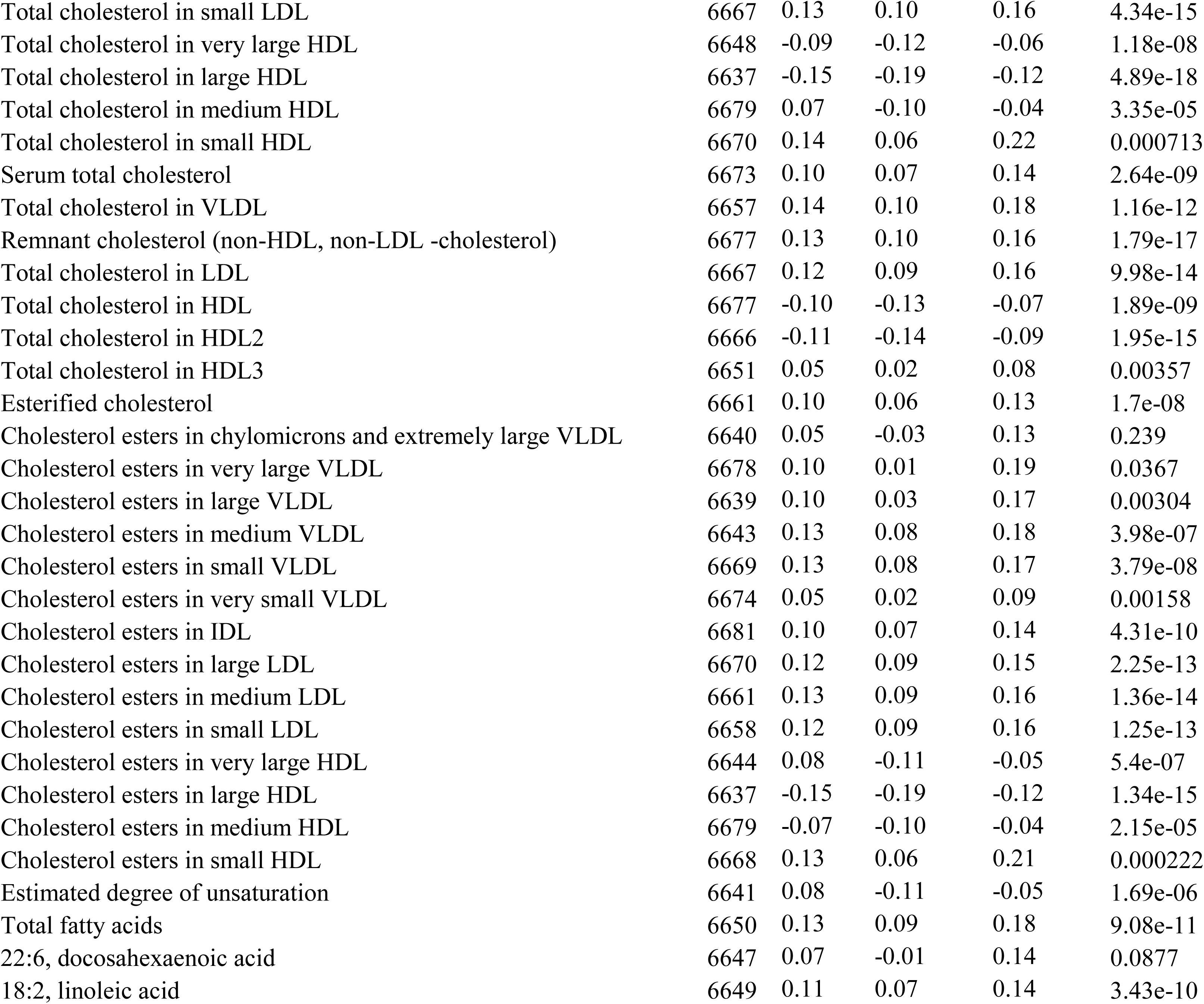
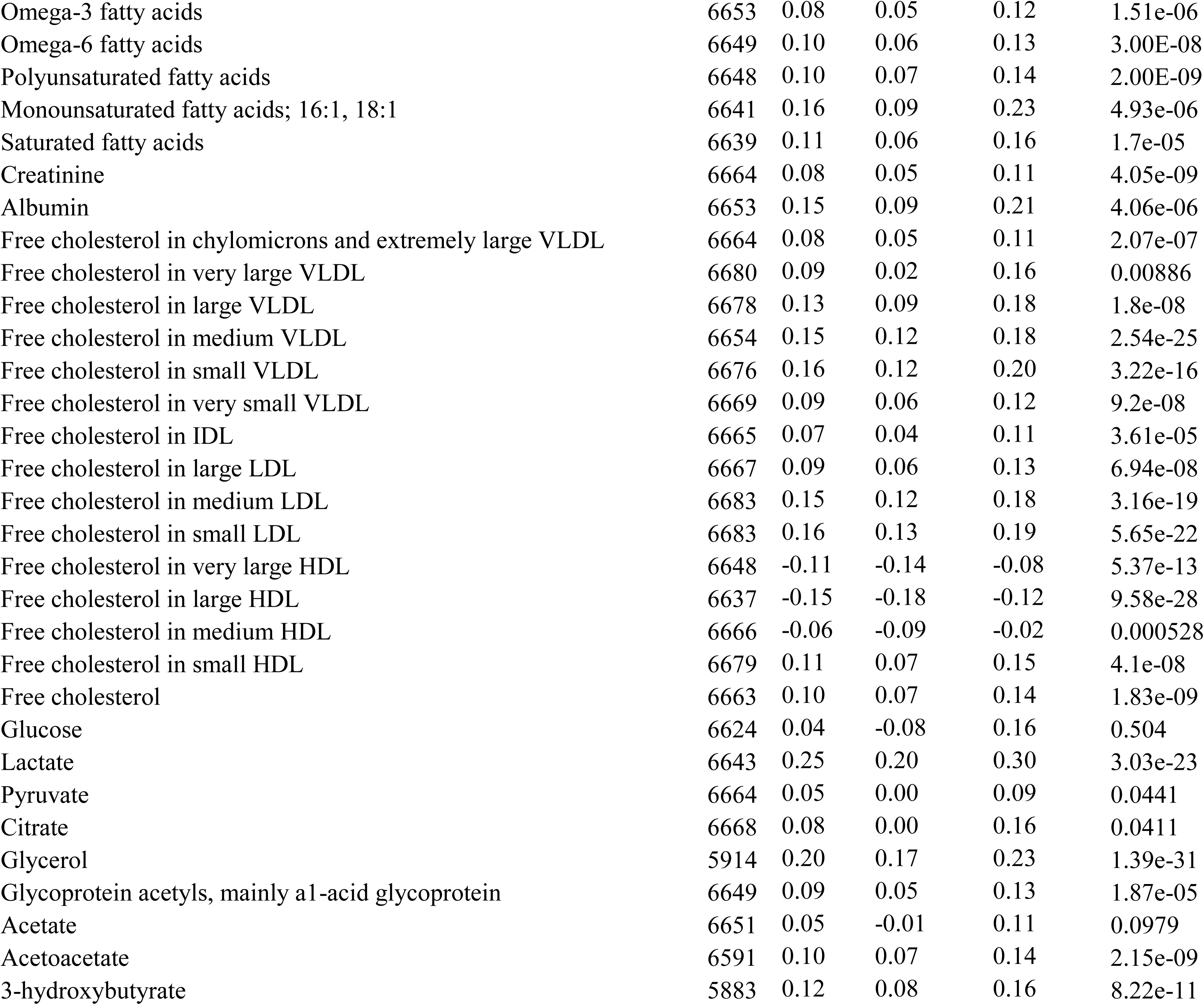
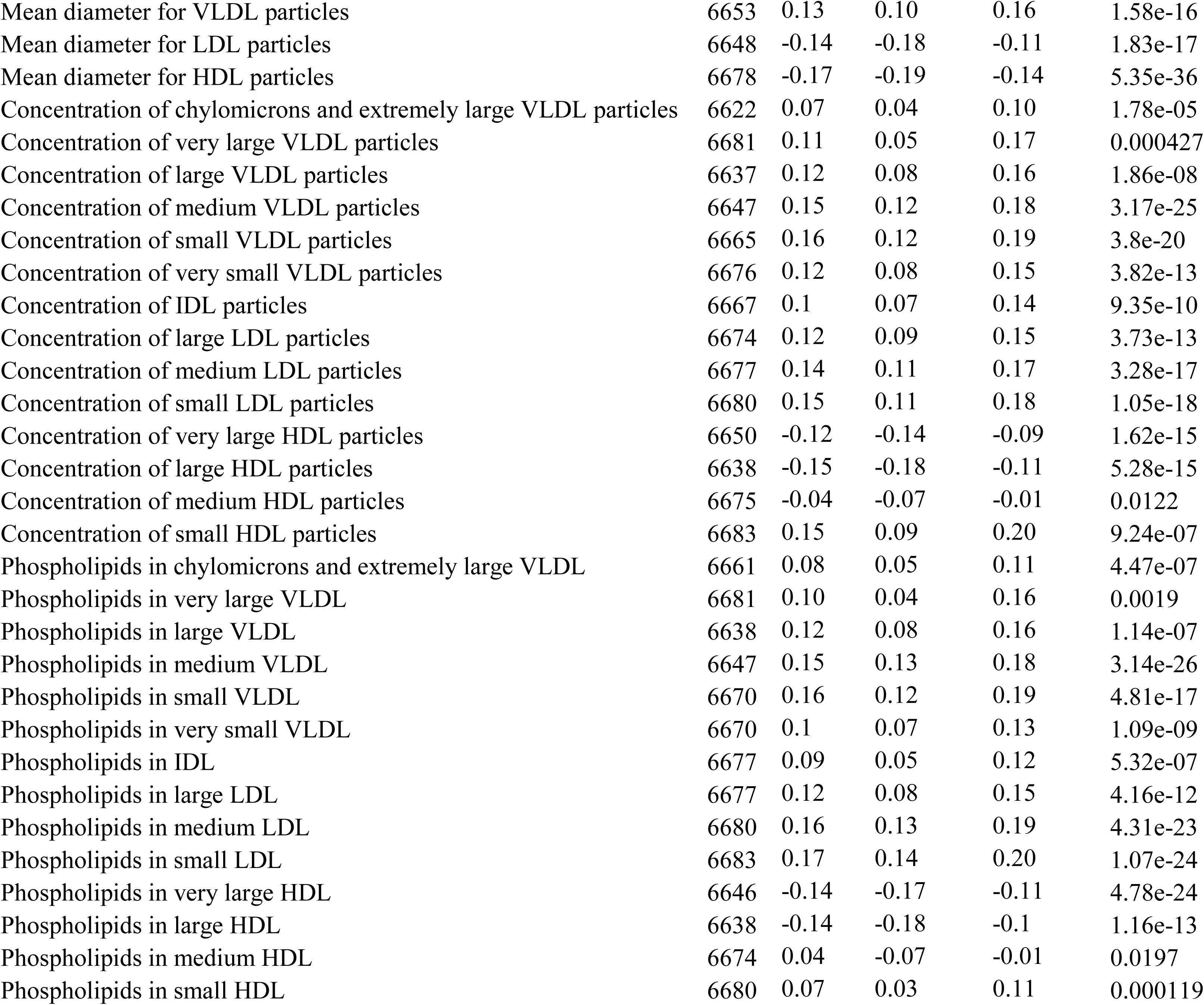
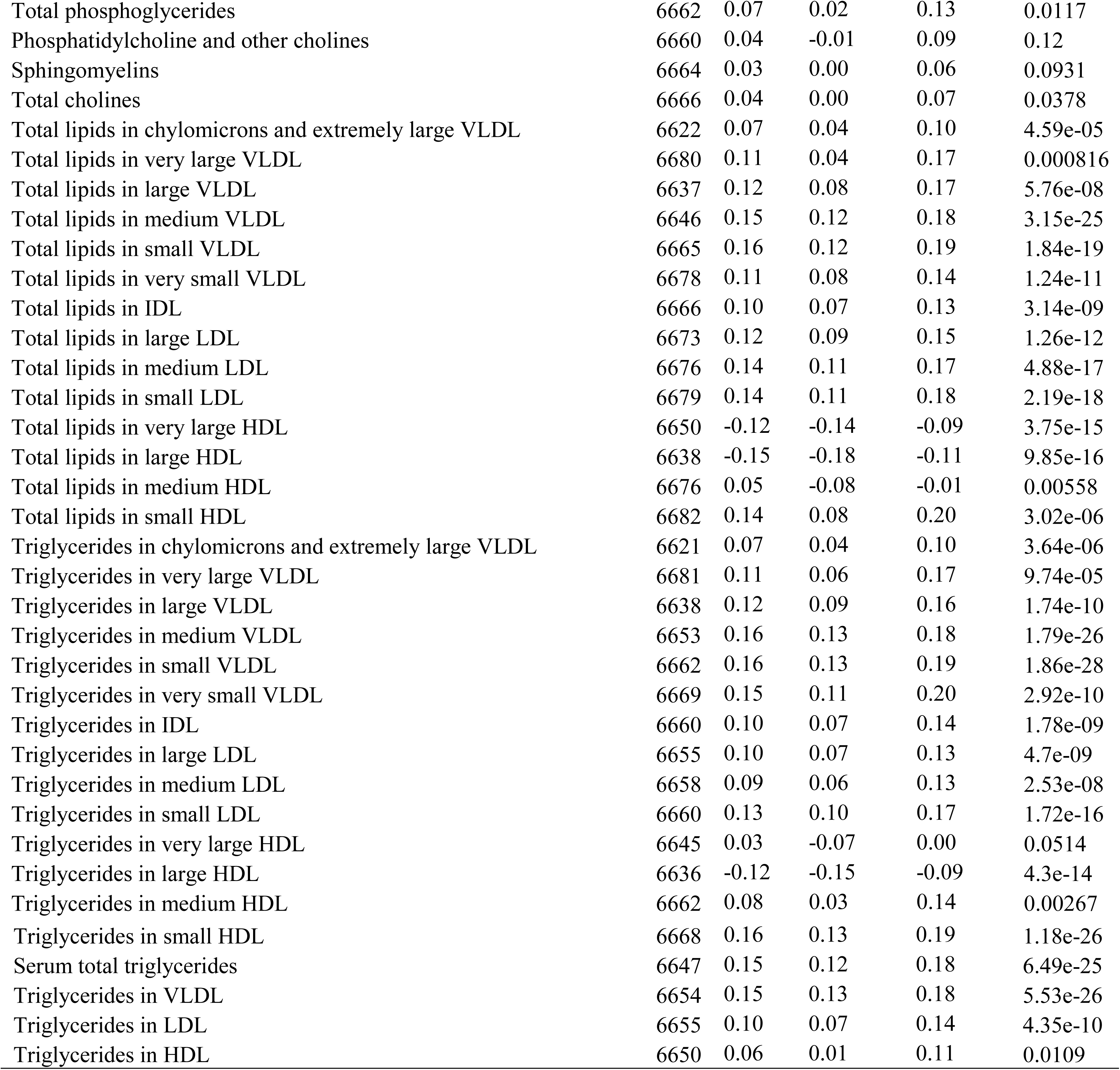
Effect sizes for association of Hb levels with metabolites. Number of participants (N) in the statistical analyses, effect sizes (Beta) in units of 1-SD change in the indicated parameter by 1-SD change in Hb, lower and upper 95% confidence intervals (CIL, CIU) and *P* values of the meta-analysis of NFBC1966 at age of 46 and YFS at age of 42. The values were adjusted for BMI. Values with ± 3 SD exclusions are shown. Users of lipid lowering medication (n = 679) and combined oral contraceptives or hormone replacement therapy (n = 391) were removed from the analyses.

**Extended data Table 9.** Effect sizes for association between Hb levels, oxygen consumption at rest and body fat percentage in a subcohort of NFBC1966 at age 31. Number of participants (N) in the statistical analyses, effect sizes (Beta) in 1-SD unit changes, lower and upper 95% confidence intervals (CIL, CIU) and *P* values and of the study population in analyses. All effects were adjusted for sex.

| Effect of | N | Beta | CIL | CIU | P |
| --- | --- | --- | --- | --- | --- |
| Hb level on oxygen consumption at rest | 114 | 0.26 | 0.04 | 0.48 | 0.02 |
| Oxygen consumption at rest on body fat percentage | 121 | 0.19 | 0.08 | 0.31 | 0.001 |
| Hb level on body fat percentage | 117 | 0.14 | 0.00 | 0.29 | 0.057 |

**Extended data Table 10.**
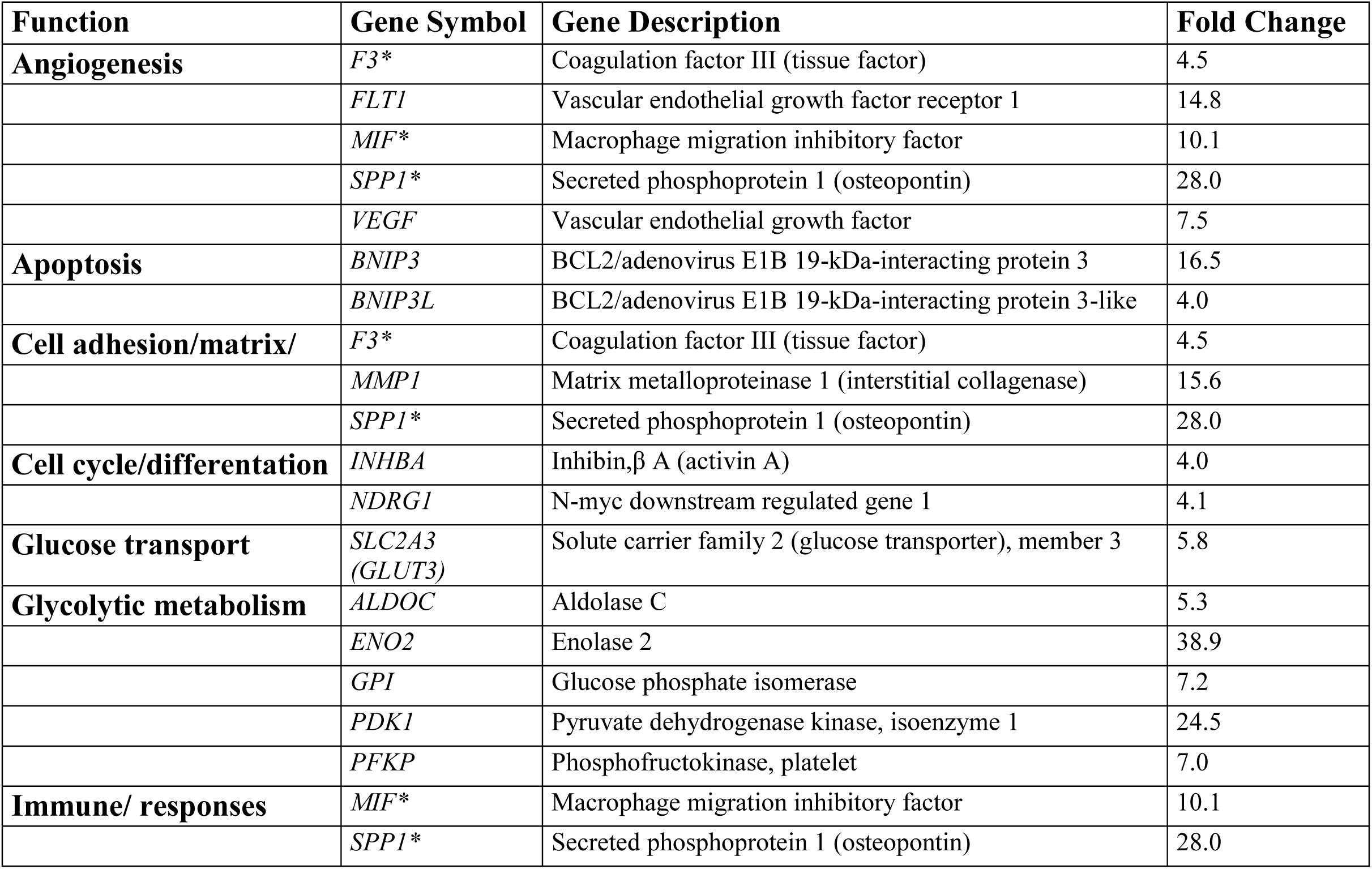
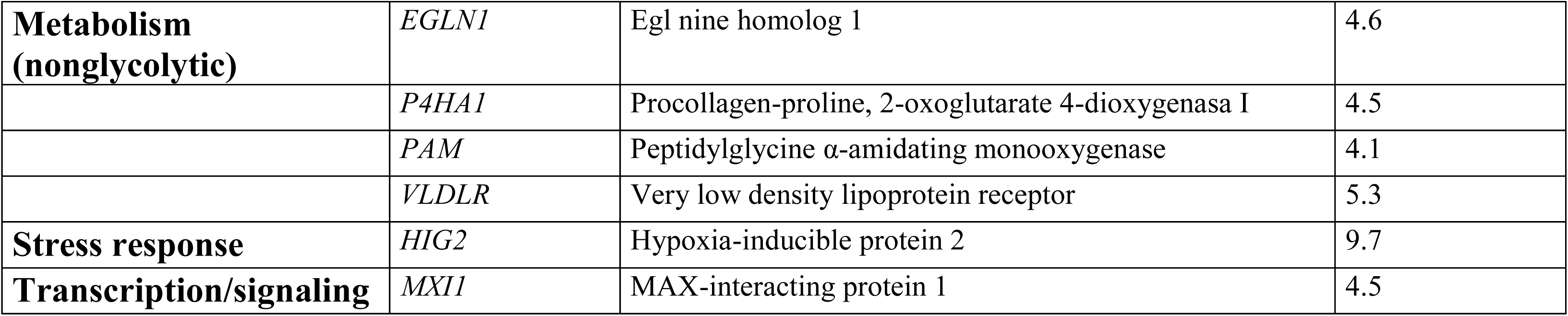
Hypoxia-induced genes used in the gene set enrichment analysis (GSEA). The genes analysed included those known to be induced at least four-fold by hypoxia in human peripheral blood monocytes (Table modified from Bosco MC et al, J Immunol. 2006, ref. 23 in the main text). Genes appearing in multiple functional categories are indicated by an asterisk.

**Extended data Table 11.**
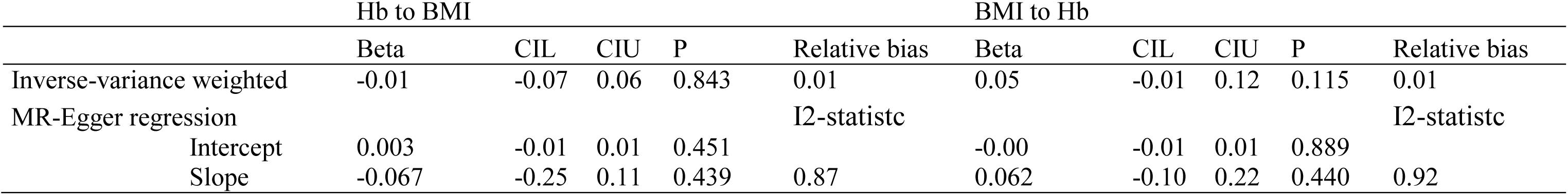
Causal estimates of association between Hb levels and BMI and *vice versa* using Mendelian randomization in European ancestry summary statistics from genome-wide meta-analysis studies. Effect sizes (Beta) in SD units, standard error (SE), lower and upper 95% confidence intervals (CIL, CIU) and *P* values of summarized data from genome-wide association analysis.

**Extended data Table 12.**
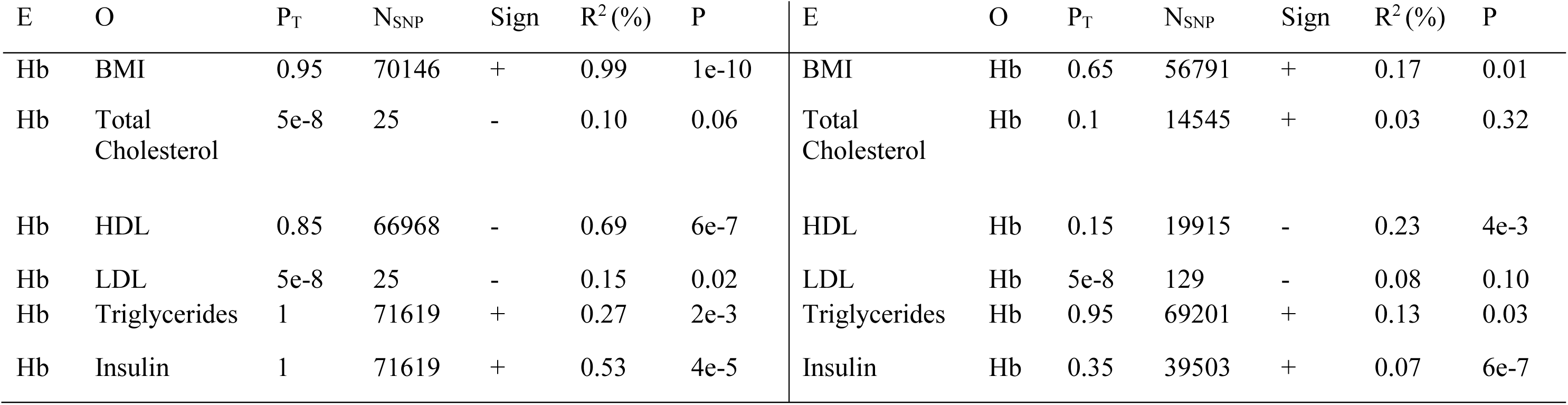
Bidirectional polygenic risk score (PRS) results. The PRS was calculated using exposure phenotype (E) GWAS results, optimised for each outcome phenotype (O) in NFBC1966 at 46 years. P_T_ = *P* value threshold for the optimised PRS. N_SNP_ = number of SNPs in the optimised PRS. *R*^*2*^ = variance explained in the outcome by the optimised PRS. P = *P* value for testing the null *R*^*2*^ = 0.

**Extended data Figure 1.**
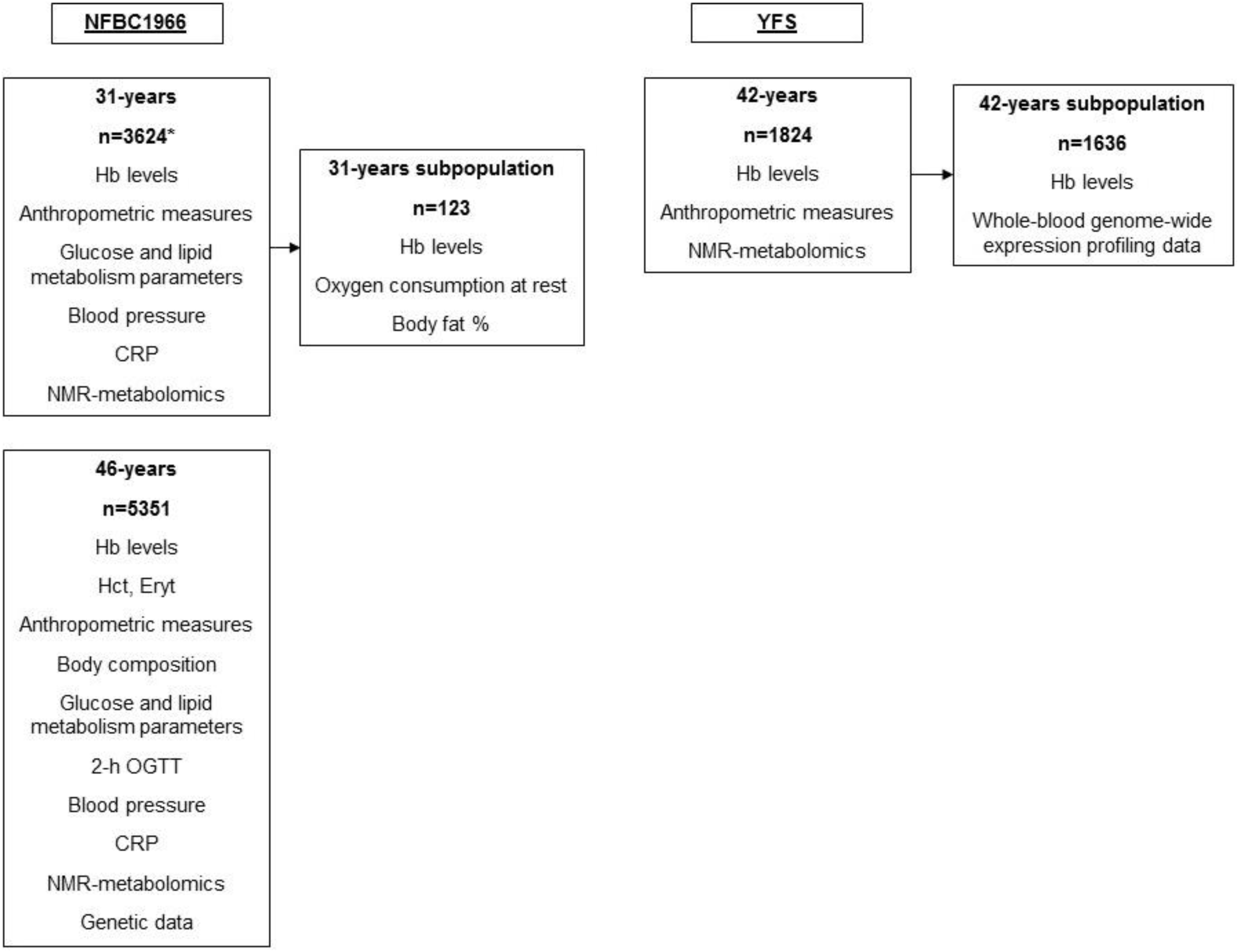
Flow chart of the study population.

**Extended data Figure 2.**
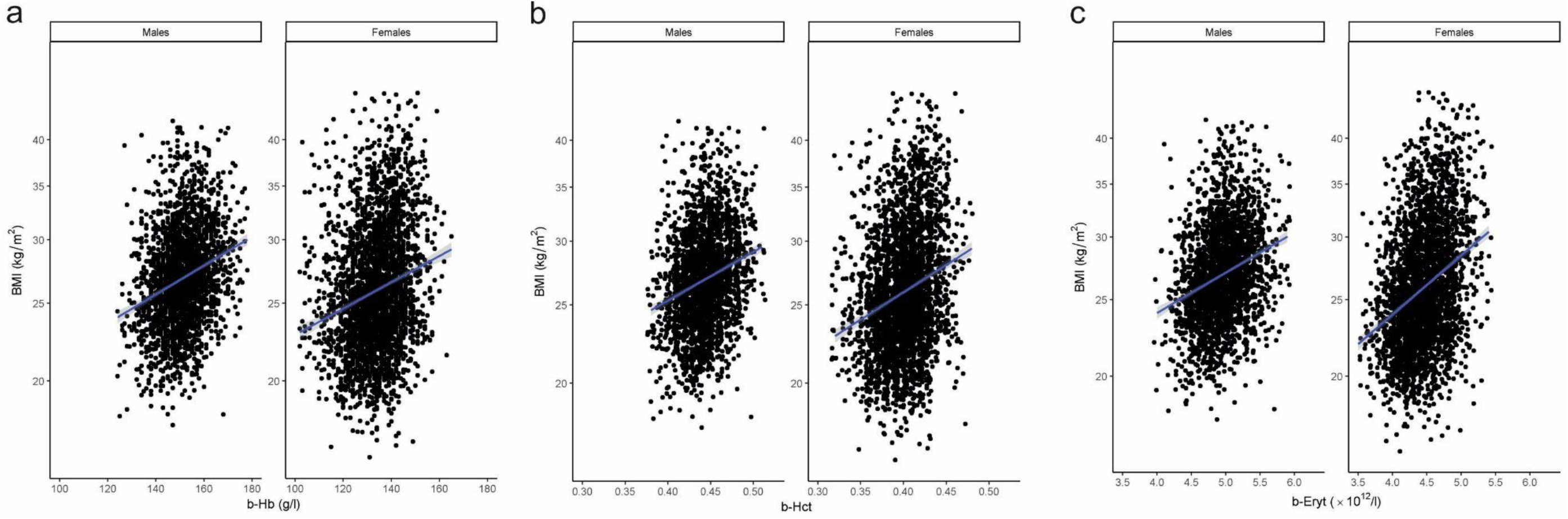
Red blood cell parameters associate with BMI in both sexes. **a**, Association of Hb levels with BMI in both genders in NFBC1966 at age of 46. **b,** Association of hematocrit levels with BMI in NFBC1966 at age of 46. **c,** Association of erythrocyte counts with BMI in NFBC1966 at age of 46. The data was adjusted smoking and physical activity. Unadjusted regression line (blue) with 95% confidence intervals (gray) are shown.

**Extended data Figure 3.**
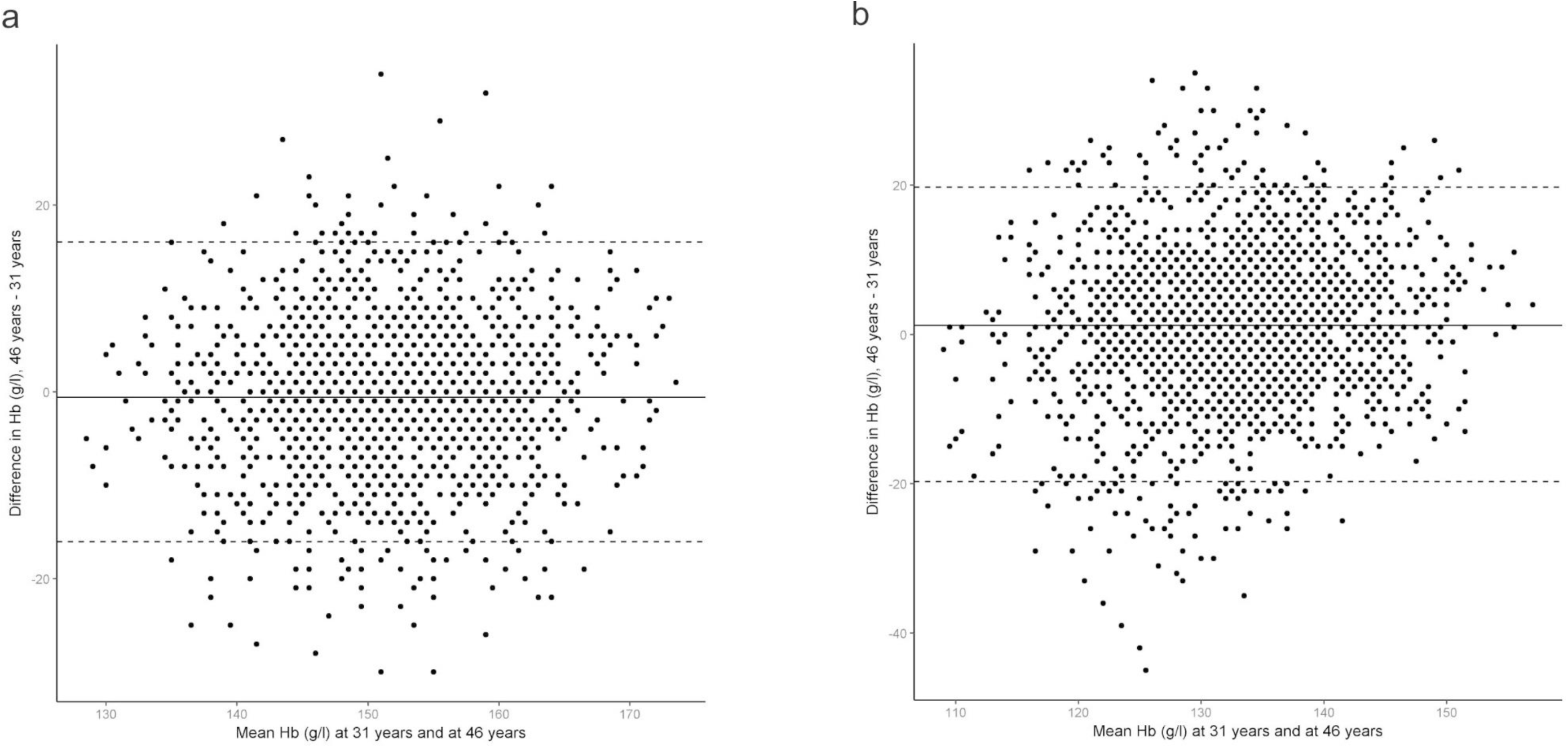
Bland-Altman plot of Hb levels at age 31 and 46 in both sexes in NFBC1966. **a,** males and **b,** females.

**Extended data Figure 4.**
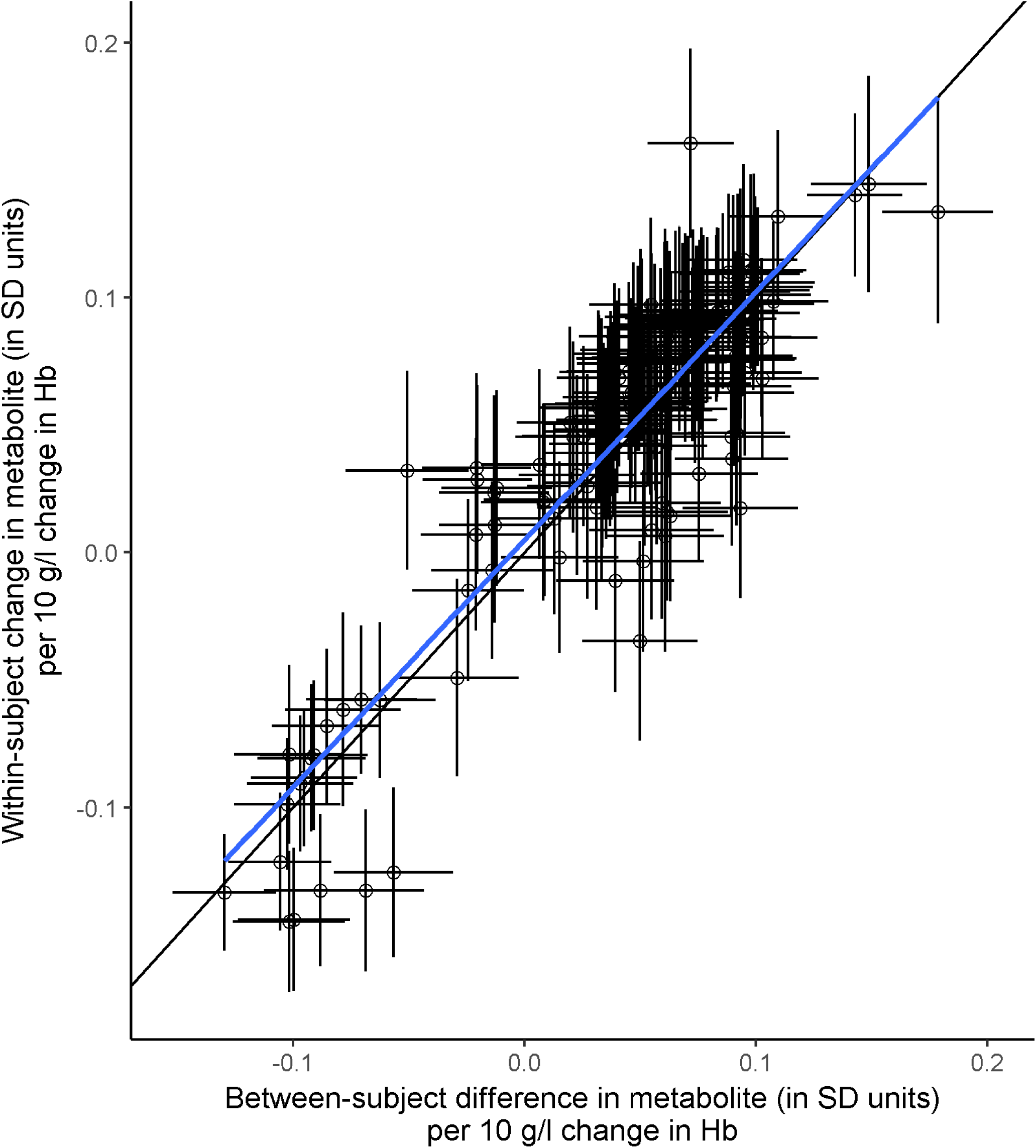
Between subject difference in Hb levels and metabolites of NFBC1966 at age of 31 vs age of 46.

**Extended data Figure 5.**
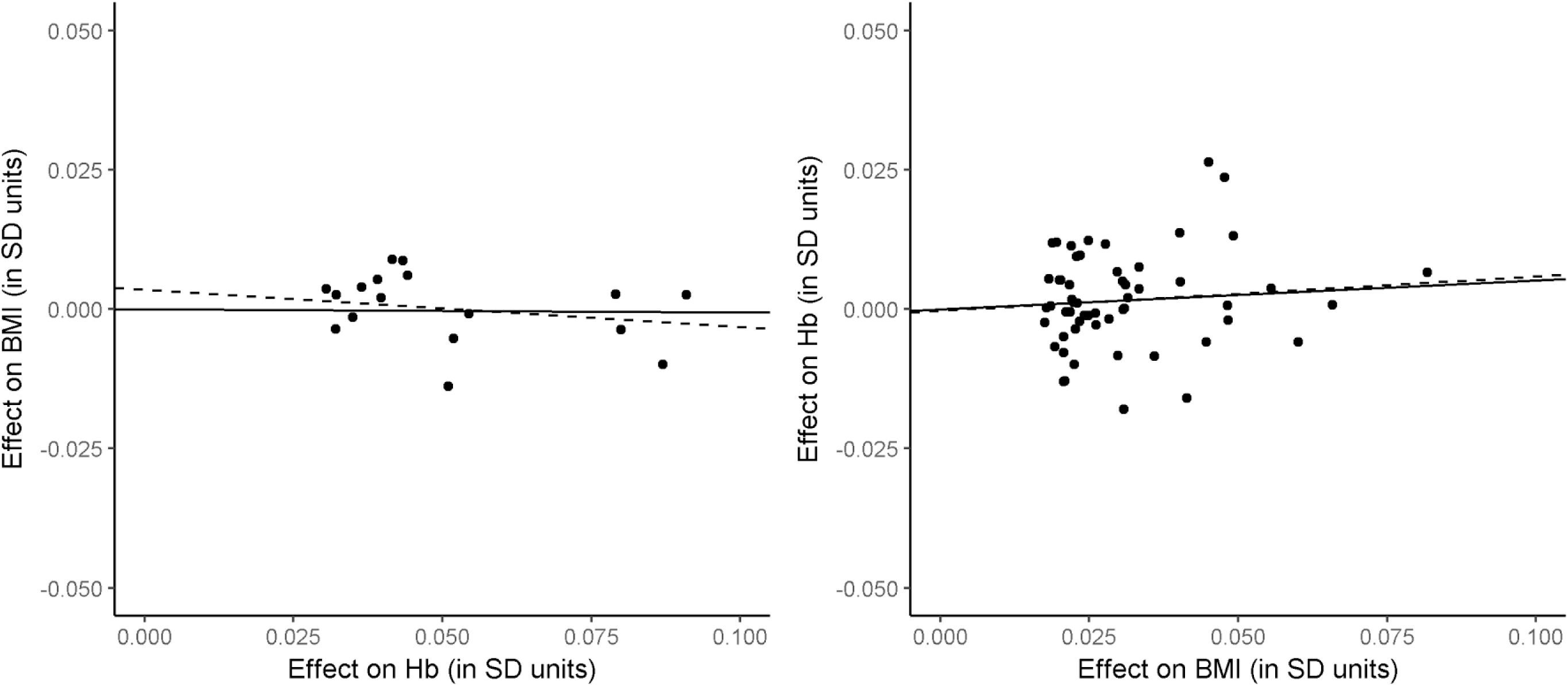
Scatter plots of SNP effects for causal effects of Hb on BMI (left) and BMI on Hb (right). The slopes of the solid line and dashed line give estimates for the causal effect using IVW method and MR-Egger regression, respectively. The intercept of the dashed line at × = 0 gives the MR-Egger regression estimate for pleiotropic effect.

**Extended data Figure 6.**
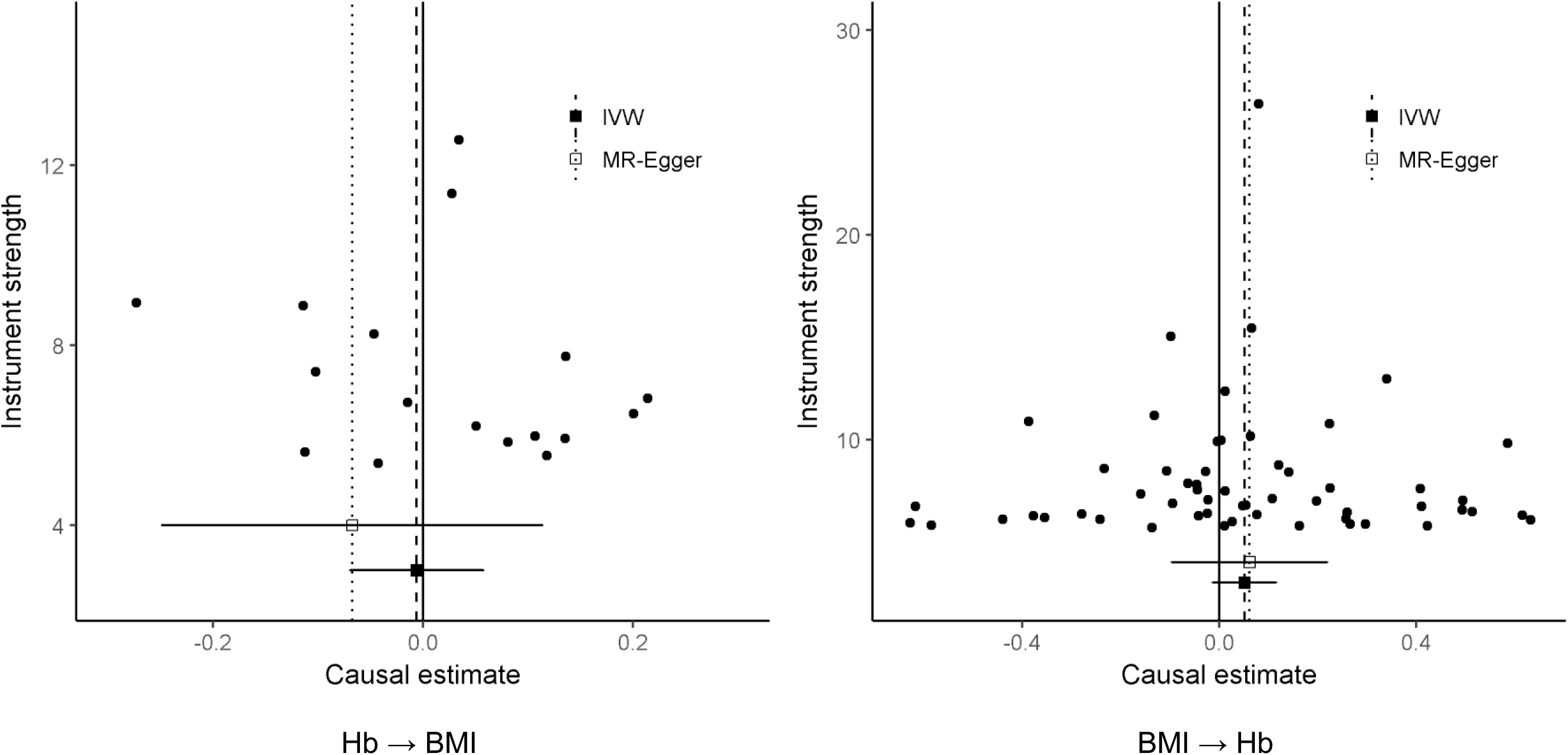
Funnel plots of effects of Hb on BMI and BMI on Hb.

**Extended data Figure 7.**
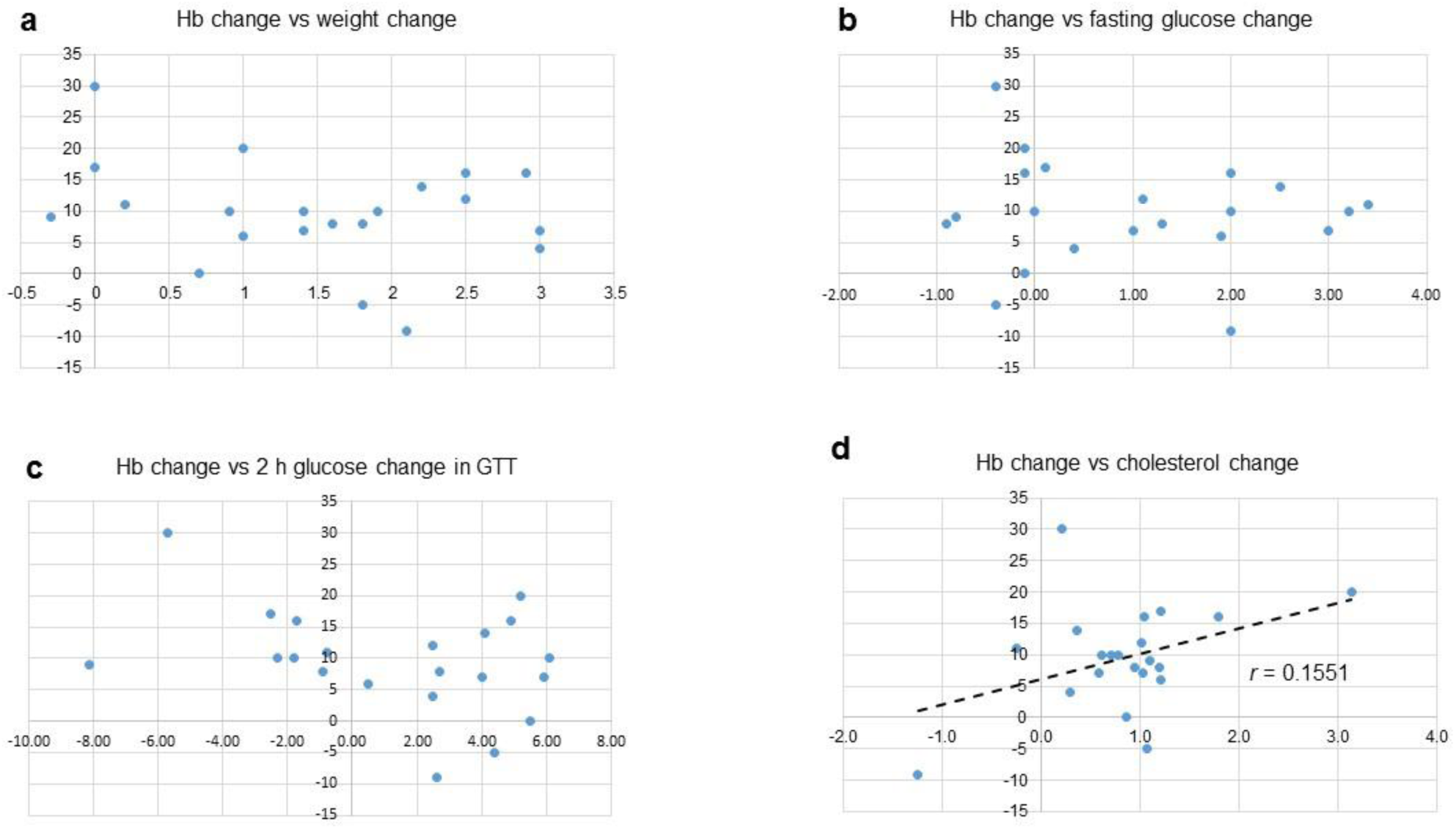
Correlation of change in Hb levels with changes in body weight, fasting blood glucose and 2 h glucose levels in GTT and serum total cholesterol levels, respectively, in individual mice following venesection. The values were determined at baseline and two weeks after venesection. The changes in them were correlated. While only the data in **d** indicate linear dose-response all data (**a**-**c**) indicate a positive response with the increase in Hb correlating with increase in body weight (**a**), higher fasting glucose levels (**b**) and 2 h glucose levels in GTT (**c**) and higher serum total cholesterol levels (**d**), respectively.

